# T- and L-type calcium channels maintain calcium oscillations in the murine zona glomerulosa

**DOI:** 10.1101/2023.06.09.544326

**Authors:** Hoang An Dinh, Marina Volkert, Ali Kerim Secener, Ute I. Scholl, Gabriel Stölting

## Abstract

The zona glomerulosa of the adrenal gland is responsible for the synthesis and release of the mineralocorticoid aldosterone. This steroid hormone regulates salt reabsorption in the kidney and blood pressure. The most important stimuli of aldosterone synthesis are the serum concentrations of angiotensin II and potassium. In response to these stimuli, voltage and intracellular calcium levels in the zona glomerulosa oscillate, providing the signal for aldosterone synthesis. It was proposed that the voltage-gated T-type calcium channel Ca_V_3.2 is necessary for the generation of these oscillations. However, *Cacna1h* knockout mice have normal plasma aldosterone levels, suggesting additional calcium entry pathways. We used a combination of calcium imaging, patch clamp and RNA sequencing to investigate such pathways in the murine zona glomerulosa. *Cacna1h*^-/-^ glomerulosa cells still showed calcium oscillations with similar concentrations as wild-type mice. No calcium channels or transporters were upregulated to compensate for the loss of Ca_V_3.2. The calcium oscillations observed were instead dependent on L-type voltage-gated calcium channels. Furthermore, we found that L-type can also partially compensate for an acute inhibition of Ca_V_3.2 in wild-type mice. Only inhibition of both, T- and L-type calcium channels abolished the increase of intracellular calcium caused by angiotensin II in wild-type. Our study demonstrates that T-type calcium channels are not strictly required to maintain glomerulosa calcium oscillations and aldosterone production and pharmacological inhibition of T-type channels alone will likely not significantly impact aldosterone production over time.

## Introduction

The adrenal glands are a pair of endocrine organs located above the kidneys. Within its cortex, steroid hormones are synthesized from the precursor cholesterol. The outermost zona glomerulosa (ZG) produces aldosterone, the zona fasciculata either cortisol in humans or corticosterone in mice and the innermost zona reticularis (in humans) androgens. Aldosterone is responsible for the maintenance of blood salt levels and volume via the regulation of various ion transporters in the kidney and intestine. Its synthesis is tightly controlled by stimuli linked to these targets, primarily the serum concentrations of potassium and angiotensin II (Ang II)^1^. ZG cells have a highly negative resting membrane potential at rest. Binding of Ang II to its cognate receptor leads to the closure of background potassium channels^2^. This results in oscillatory depolarizations of ZG cells^3^, causing similarly oscillatory influx of calcium via voltage-gated calcium channels (VGCCs) ^2,4–9^.

Calcium itself is required for several key functions in ZG cells, such as the regulation of transcription factors upstream of the aldosterone synthase *Cyp11b2*^10,11^ and the transport of cholesterol to the inner membrane leaflet of mitochondria for conversion into aldosterone^12^. Tight control over [Ca^2+^]_int_ is therefore key to the control of aldosterone synthesis.

The original publication identifying murine ZG cells as voltage oscillators demonstrated that voltage fluctuations critically depend on the function of the T-type VGCC Ca_V_3.2 (Gene: *Cacna1h*) and specific inhibition of this channel abolished all voltage oscillations^3^. Even before, the specific inhibition of T-type VGCCs has been proposed as a potential strategy to pharmacologically suppress aldosterone production^13^.

However, despite its critical role in the electrical excitability of the ZG, Ca_V_3.2 knock-out mice did not show altered systemic aldosterone or renin levels^8,14^. The molecular mechanisms that sustain aldosterone production in Ca_V_3.2 knock-out mice currently remain unclear. We here set out to investigate the regulation of [Ca^2+^]_int_ in the ZG of mice lacking Ca_V_3.2 and the implications of the results on wild-type (WT) ZG function and pharmacology.

## Methods

Detailed methods are available in the supplemental material.

### Mice and organ harvest

WT and Cacna1h KO mice^8^ were kept in IVF cages with water and food ad libitum according to local regulations. For extraction of adrenal glands, mice were anesthetized using isoflurane and euthanized by cervical dislocation.

### Calcium imaging

Calbryte 520 AM- or Fura-2 AM-stained acute slices (120 µm thickness) were constantly perfused with bicarbonate-buffered, oxygenated solution. K^+^, Ang II and inhibitors were added in indicated concentrations.

### Electrophysiological recordings of dissociated adrenal cortical cells

The adrenal cortex was manually dissected and cells isolated by enzyme- and shear-based dissociation. Cells were used for whole-cell patch clamp recordings the next day.

### Bulk RNA-seq

RNA was extracted from dissected whole adrenal cortices (6 WT and 6 KO mice; 3 male, 3 females each). RNA was processed and sequenced in the Genomics core facility of the Berlin Institute of Health.

### Single nucleus RNA-seq

Adrenal glands (two males and two females) were processed to obtain a nuclei suspension. Sequencing-ready libraries were generated from these and sequencing was performed on a HiSeq 4000 device.

### Statistics

All P-values (except for the differential expression analysis) are results from a likelihood ratio test of linear mixed models and are indicated as follows: ns, P ≥ 0.05; *, <0.05; **, P<0.01; ***, P<0.001. All error bars and bands show the 95% confidence interval of the mean value.

For differential expression analysis, P-values were adjusted by Benjamini-Hochberg for multiple comparisons

## Results

### ZG cells from *Cacna1h*^-/-^ mice still exhibit intracellular calcium oscillations

To study [Ca^2+^]_int_ in murine ZG cells, we stained acutely prepared slices from mouse adrenal glands with the fluorescent calcium indicators Calbryte 520 AM (relative signal but high temporal resolution) or Fura-2 AM (allows for [Ca^2+^]_int_ quantification). We used WT as well as *Cacna1h*^*-/-*^ (KO) mouse lines that we established previously^8^.

When stimulated with 500 pmol/l Ang II and 4 mmol/l K^+^, ZG cells in slices from WT mice responded with pronounced Ca^2+^ oscillations. As observed previously^7,8^, individual transients (spikes) were homogeneous in appearance. Spikes were clustered in bursts, separated by pauses at a mostly constant baseline [Ca^2+^]_int_ (Fig. 1A). Although lacking the Ca_V_3.2 channel, [Ca^2+^]_int_ in ZG cells from KO mice exhibited similar behavior: Homogeneous spikes, clustered into bursts (Fig. 1B).

**Figure 1.**
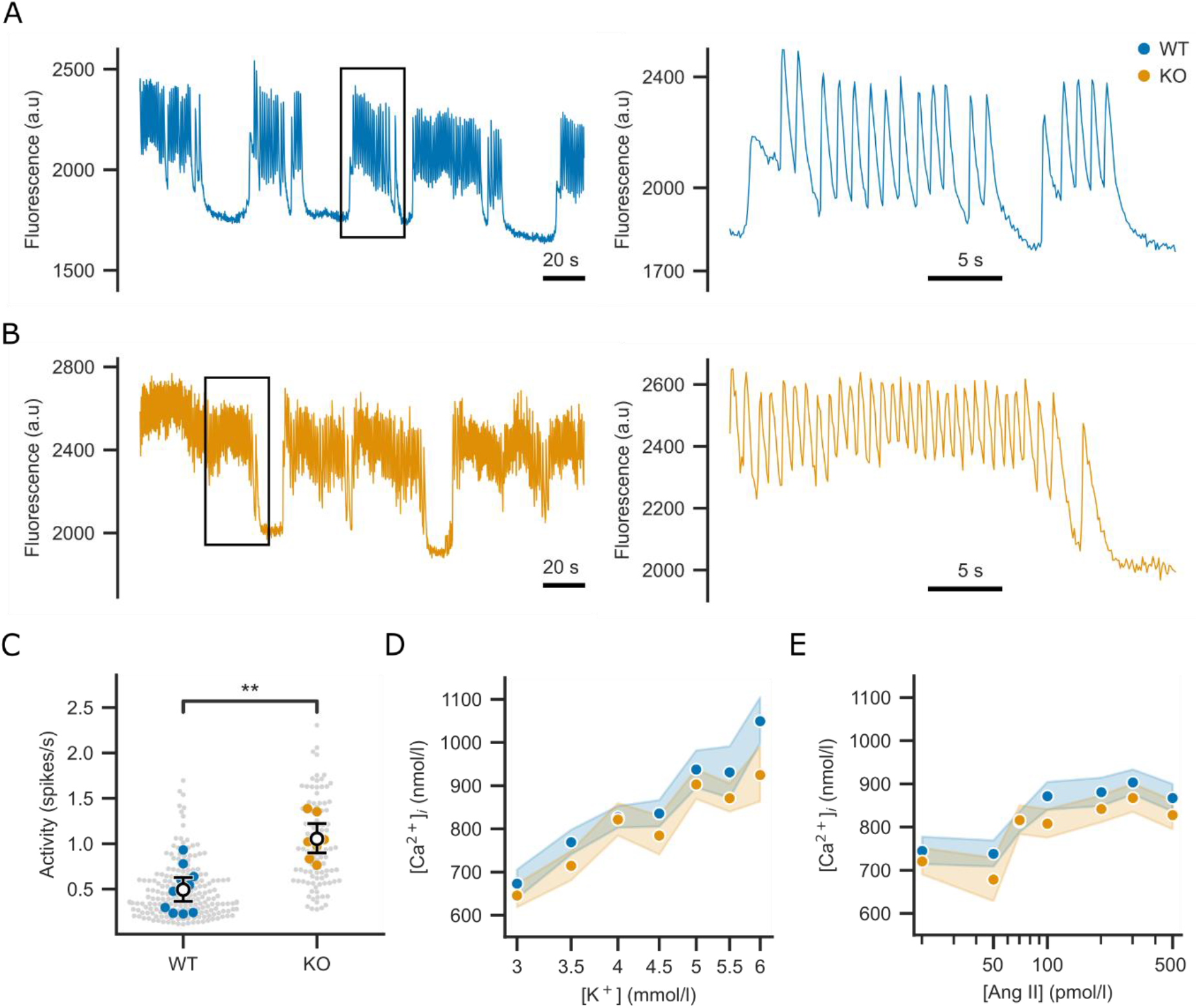
Calcium oscillations persist in *Cacna1h*^-/-^ mice. **(A-B)** Fluorescence signals recorded from one representative ZG cell each of a WT **(A)** or KO **(B)** Calbryte 520 AM-stained adrenal slice and stimulated with 4 mmol/l K^+^ and 500 pmol/l Ang II. A 30 s magnification (indicated by the black rectangle) is shown on the right. **(C)** Calcium spike activity recorded in ZG cells from Calbryte 520 AM-stained adrenal slices over 7.5 min is lower in WT than in KO **(D-E)** Mean [Ca^2+^]_int_ in ZG cells of Fura-2 AM-stained adrenal slices. The Ang II concentration was kept constant at 100 pmol/l in **(D)** while potassium was fixed at 4 mmol/l in **(E)**.

The most prominent difference of the spiking in KO mice was the significantly increased overall number of spikes per second (activity; Fig. 1C, Supplementary Table S1). This was caused by an increased frequency of spikes within bursts and shorter gaps between bursts rather than longer bursts (Supplementary Fig. S1A-C).

### ZG stimulation results in similar changes of intracellular calcium levels in KO and WT mice

We also investigated mean [Ca^2+^]_int_ over a wide range of concentrations of potassium (Fig. 1D) and Ang II (Fig. 1E). With increasing stimulation, [Ca^2+^]_int_ similarly increased in ZG cells of both, WT and KO. Mean [Ca^2+^]_int_ were not significantly different between genotypes (Supplementary Tables S2 and S3), suggesting that the higher activity and intra-burst frequency of calcium spikes in KO cells does not result in higher calcium influx over time but rather maintains the sensitivity to physiological stimuli of aldosterone production.

### Bulk RNA-seq of adrenal cortices from KO mice revealed no upregulation of calcium transport genes

To identify differentially expressed genes between genotypes and to quantify the expression of VGCCs in WT, we performed bulk RNA-sequencing of adrenal cortices from 6 WT and 5 KO mice.

In the WT adrenal cortex, *Cacna1h* (Ca_V_3.2), *Cacna1c* (Ca_V_1.2) and *Cacna1d* (Ca_V_1.3) were the most strongly expressed VGCC α1-subunit genes (Fig. 2A, Supplementary Table S4). The most prominent subunits were β2 and α2δ1 (Supplementary Table S4).

**Figure 2.**
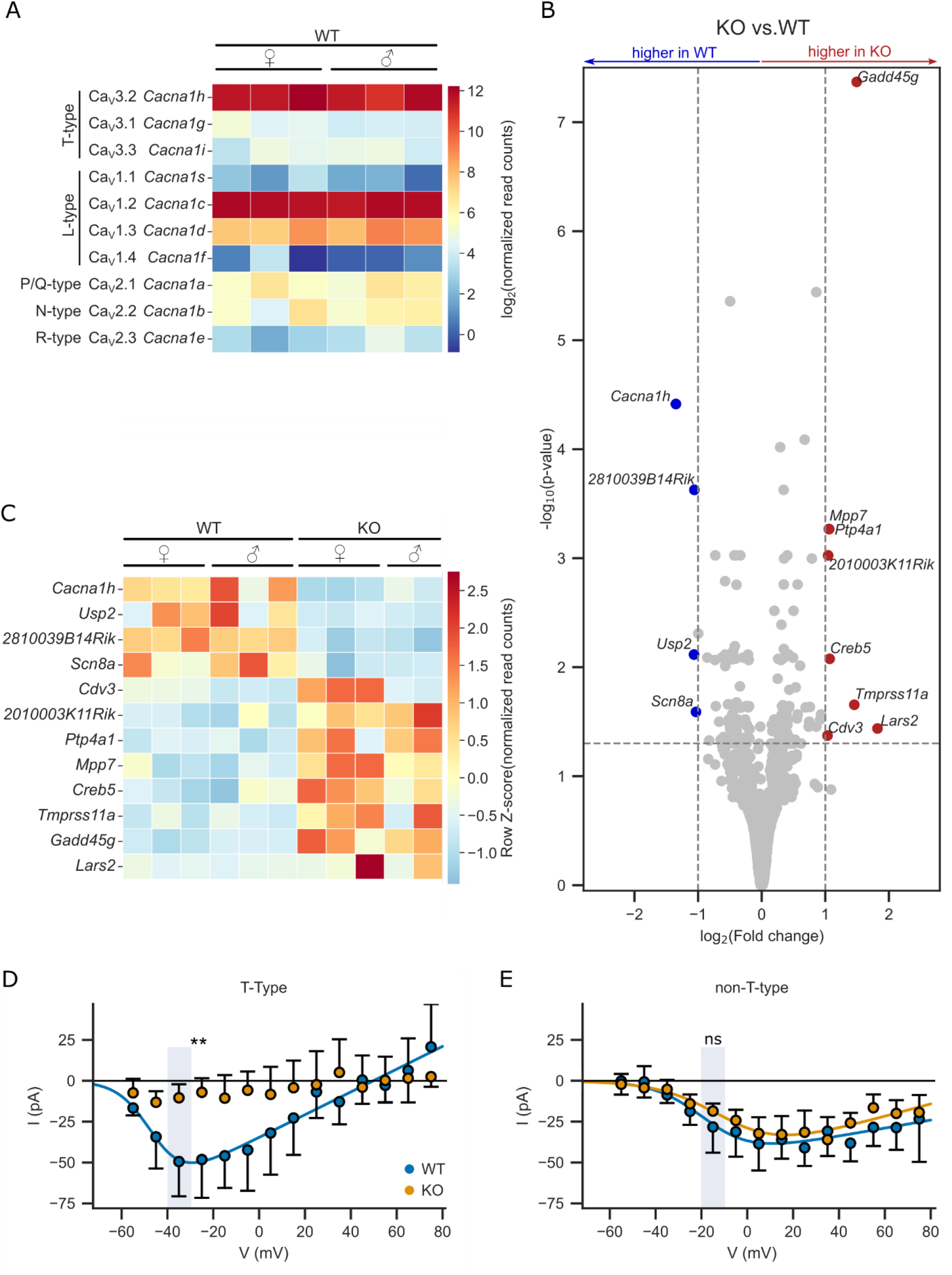
RNA-seq reveals only few differentially expressed genes. **(A)** Heat map illustrating the normalized log_2_ transformed read counts of VGCC transcripts detected in WT samples (columns, n_animals,WT_ = 6). **(B)** Volcano plots with the log_2_-fold change (LFC, cutoff = |1|) in gene expression in KO vs. WT samples (n_animals,WT_ = 6, n_animals,KO_ = 5) and the log_10_ transformed statistical significance (p-value, cutoff=0.05). The 12 differentially expressed genes (DEGs) are highlighted as up- (red) or down (blue) regulated. **(C)** Heatmap illustrating the row Z-score of the normalized read counts of DEGs sorted by their LFC. **(D-E)** Isolated adrenocortical cells from KO mice lack T-type currents as shown by whole cell patch-clamp recordings. Voltage dependence of the T-type **(D)** and non-T-type **(E)** peak current amplitudes. Solid lines represent fits of equation 1 to our experimental data (see supplementary material). Circles represent mean values per cell. Only one half of the CI is displayed.

Apart from a downregulation of *Cacna1h* mRNA, likely due to nonsense-mediated decay, we did not observe any significant changes to known calcium channel genes in KO compared to WT mice (Supplementary Table S4). In total, we observed 12 differentially expressed genes with at least a 2-fold change between genotypes (Fig. 2B and 2C, Supplementary Table S5). Of these, besides *Cacna1h*, only *Creb5* has previously been associated with regulating aldosterone synthesis^15^.

Looking at other known genes involved in aldosterone synthesis (according to the KEGG database) without cutoff for the change in expression, we found 3 significant changes in KO (Agtr1a: 1.35-fold, p_adj_ = 0.048; Star: 1.27-fold, p_adj_ = 0.007; Gna11: 0.87-fold, padj = 0.023; Supplementary Table S6). However, these changes were small and are therefore unlikely to carry a large functional impact on their own.

GO term and KEGG pathway overrepresentation analyses for the set of differentially expressed genes did not yield any significant results.

### Adrenocortical cells from KO mice do not exhibit T-type calcium currents

To investigate functional changes of existing calcium channels and to confirm the loss of Ca_V_3.2 in our KO mice, we performed whole cell patch clamp recordings from isolated adrenocortical cells^5^. Currents (Supplementary Fig. S2) were recorded in response to two separate voltage clamp protocols^16^: First from a holding potential of -80 mV, then from a holding potential of -40 mV. At -80 mV, VGCCs reside in the closed but activatable state leading to currents from all different types (Supplementary Fig. S2, left). With the second protocol (from -40 mV), the low-voltage activated T-type channels are already inactivated before the recording. Only non-T-type VGCCs (including the L-type channels Ca_V_1.2 and Ca_V_1.3) remain available (Supplementary Fig S2, middle). The subtraction of the current from the holding potential of -40 mV from the one from -80 mV revealed the T-type channel component (Supplementary Fig. S2, right). In WT cells, we observed both, T- and non-T-type currents in all recorded cells (7 cells from 3 animals, Fig. 2D-E). In KO cells, we did not observe T-type currents in any of the recorded KO cells (9 cells from 4 animals, Fig. 2D).

Non-T-type currents observed in KO cells were not different in their maximum amplitude or voltage dependence compared to WT cells (Fig. 2E, Supplementary Table S7). This excludes large post-translational modifications to increase L-type currents in response to the loss of Ca_V_3.2.

### Ca_V_3.2 is not required for calcium oscillations in WT mice

RNA-seq and patch clamp suggested Ca_V_3.2 as the only relevant T-type calcium channel in adrenocortical cells. To confirm its importance for calcium oscillations in the ZG, we studied the effect of the T-type calcium channel inhibitor TTA-P2^17^ in adrenal slice preparations using calcium imaging. At the chosen concentration of 15 µmol/l, virtually all T-but less than 10 % of L-type VGCCs^18^ are expected to be inhibited.

Calcium oscillations in ZG cells from KO mice were unaffected by the presence of TTA-P2 (Fig. 3A), confirming the functional absence of other T-type channel isoforms. Treatment of WT slices led to a heterogeneous response, with some cells exhibiting continued calcium spiking (Fig. 3B) and others turning completely silent (Fig. 3C). Overall, ∼34% of the WT cells and almost all KO cells remained active (WT: 34.4%, 95% CI: 22.2-47.3%; KO: 90.3%, 95% CI: 84.3-95.1%; Fig. 3D). This resulted in a reduction of the mean activity in WT by about 45% when compared to control recordings (Fig. 3E and F; Supplementary Table S8-9; absolute data in Supplementary Fig. S3A and Table S10-11).

**Figure 3.**
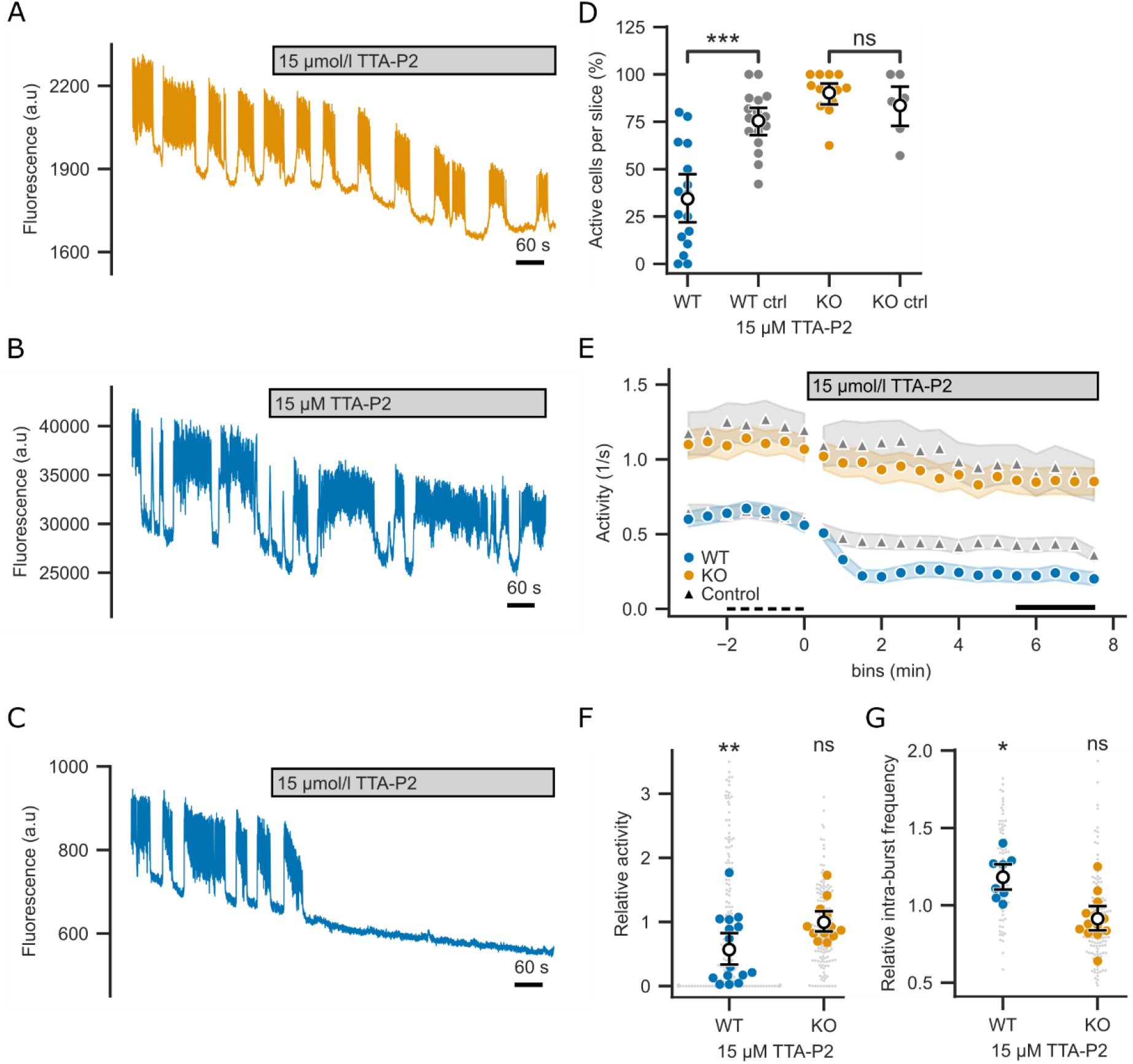
TTA-P2 only partially inhibits calcium oscillations in ZG cells of WT mice. **(A-C)** Variations in the fluorescence recorded in one representative KO **(A)** and two WT **(B-C)** ZG cells. Application of TTA-P2 inhibits calcium spikes in a fraction of the WT cells. **(D)** Fraction of cells in a slice that are still active under TTA-P2 (determined at time span 5.5 - 7.5 minutes after addition of TTA-P2, indicated by the black line in E) compared to the number of active cells before application of TTA-P2 (−2 - 0 minutes, i.e., baseline, indicated by the black dashed line in E). **(E)** In WT but not KO ZG cells, calcium spike activity was lowered by TTA-P2 when compared to controls (no TTA-P2, triangles). Relative activity **(F)** and relative intra-burst frequency **(G)** between 5.5 – 7.5 minutes after the start of the perfusion was significantly changed by TTA-P2 in WT but not KO. Data per genotype in F and G was calculated relative to and tested against control recordings. Cells were from adrenal slices stained with Calbryte 520 AM and perfused with 4 mmol/l K^+^ and 500 pmol/l Ang II.

WT cells with remaining activity in the presence of TTA-P2 responded with an increased intra-burst frequency but stayed below the level seen in KO (Fig. 3G; Supplementary Table S10-12; absolute data in Supplementary Fig. S3B, C).

These findings support an important role of Ca_V_3.2 in the generation of calcium spikes in WT mice. Still, at least in a subset of cells, other calcium influx pathways exist that can maintain calcium oscillations when T-type calcium channels are inhibited.

### L-Type calcium channels mediate calcium oscillations in KO mice

Our RNA-seq data and previous publications also demonstrated the expression of L-type ^19–21^ VGCCs in the adrenal cortex. We perfused adrenal slices with the specific L-type channel inhibitor isradipine to isolate their contribution to calcium signals in the ZG.

The concentration of isradipine (0.05 µmol/l) was chosen to be specific for L-type over other VGCCs^22–25^. The onset of inhibition was slow but resulted in an almost complete termination of calcium oscillations in KO cells. On the other hand, isradipine had nearly no effect on WT cells (Fig. 4A-C; Supplementary Fig. S4A; Supplementary Table S13-15). The intra-burst frequency remained unaffected by the inhibition of L-type channels in both genotypes (Supplementary Fig. S4B-C; Supplementary Table S14-16).

**Figure 4.**
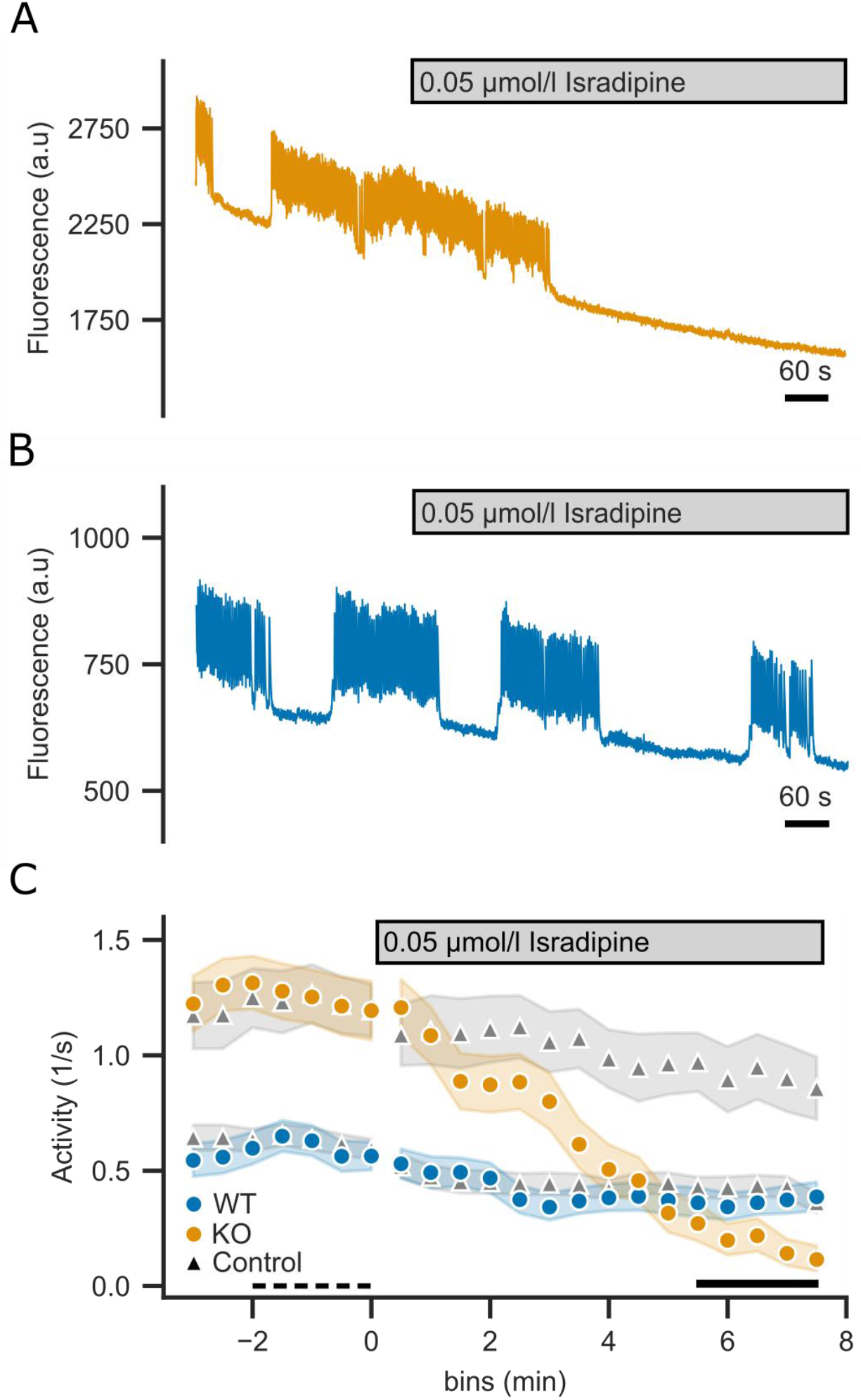
The L-type inhibitor isradipine inhibits calcium oscillations in KO but not WT cells. **(A-B)** Variations in the fluorescence recorded in one representative KO **(A)** and WT **(B)** ZG cell. Application of isradipine inhibits calcium spikes in almost all KO but not WT cells. **(C)** Calcium spike activity. Cells were from adrenal slices stained with Calbryte 520 AM and perfused with 4 mmol/l K^+^ and 500 pmol/l Ang II.

Overall, these results indicate that L-type calcium channels are essential to generate the activity in mice chronically lacking Ca_V_3.2. However, acute inhibition of L-type channels alone did not significantly change the calcium spiking in WT ZG cells.

### Both, T- and L-type calcium channels are needed for Ang II-dependent calcium signaling

To test whether L-type VGCCs underly the remaining activity in WT cells subjected to TTA-P2 or other (non T-nor L-type) VGCCs contribute to the generation of [Ca^2+^]_int_ oscillations, we studied the situation in ZG cells upon inhibition using both, TTA-P2 and isradipine. The protocol consisted of a 10-minute period in which the slice was perfused with the T-type VGCC inhibitor TTA-P2 alone, followed by another 10 minutes during which TTA-P2 was supplemented with isradipine to additionally block L-type VGCCs. To maximize the specificity of the inhibition of T-over L-type channels, we lowered the concentration of TTA-P2 to 5 µmol/l. Furthermore, we increased the isradipine concentration to 300 nmol/l to fully inhibit Ca_V_1.3^22^. A potential unspecific inhibition of T-type channels was considered less important in this context given the parallel incubation with TTA-P2. (Fig. 5A, B).

**Figure 5.**
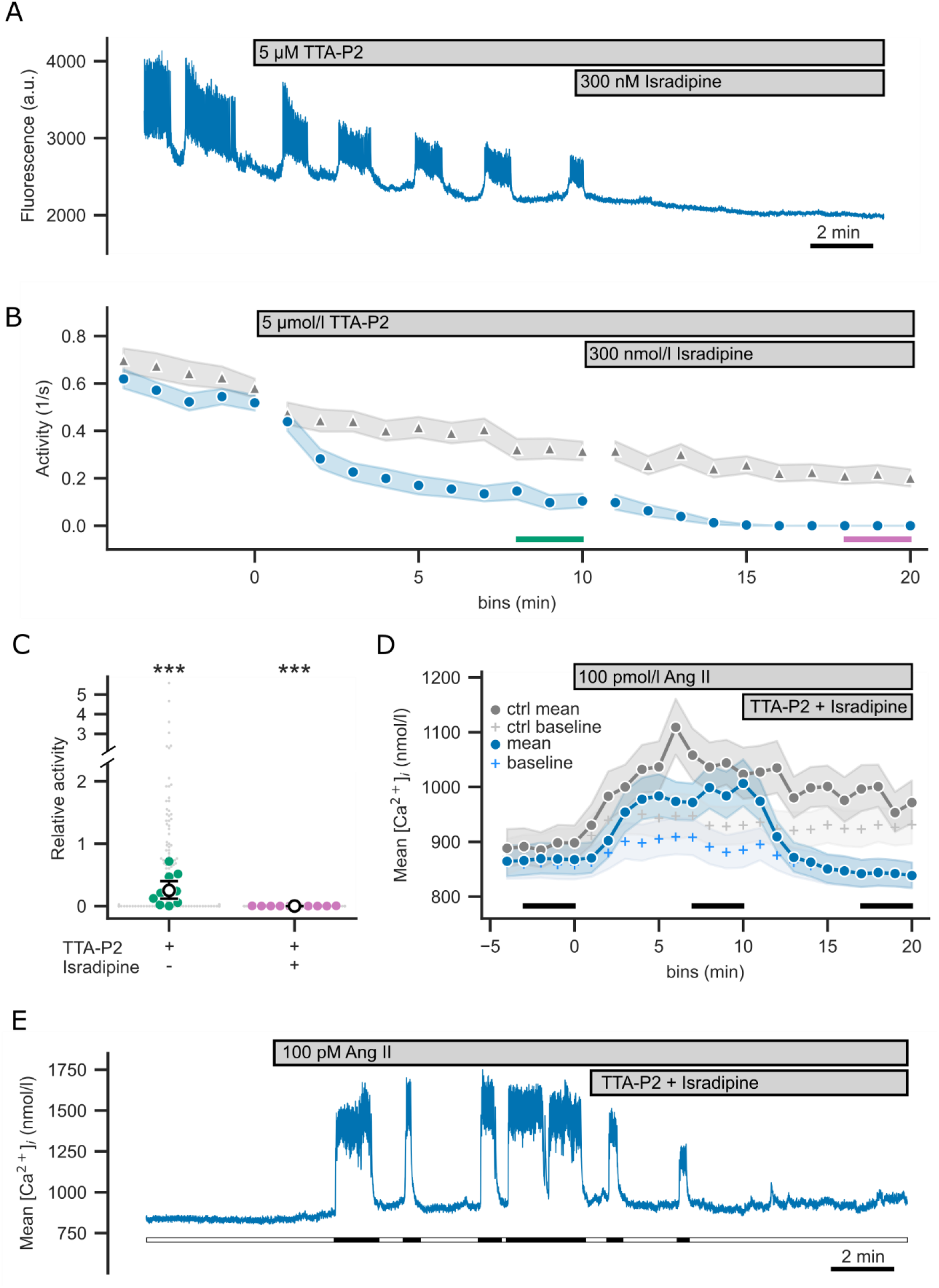
Simultaneous inhibition of L- and T-type calcium channels fully abolishes calcium signaling. **(A)** Fluorescence intensity recorded from one representative WT ZG cell. **(B)** Calcium spike activity of WT ZG cells was lowered by TTA-P2 and isradipine compared to control (no blockers, grey triangle). The last two minutes of each condition of the recording are highlighted with color-coded lines. **(C)** Calcium activity in ZG cells from WT during the steady-state phases as highlighted in B relative to and tested against controls. **(D)** Mean and baseline [Ca^2+^]_i_ in Fura-2 AM-stained ZG cells. Ang II-stimulated increase in [Ca^2+^]_i_ was abolished by inhibitors (grey: control, no inhibitors). **(E)** Representative trace of [Ca^2+^]_i_ of a ZG WT cell (unfilled bars: baseline). Cells in A-C were from adrenal slices stained with Calbryte 520 AM and perfused with 4 mmol/l K^+^ and 500 pmol/l Ang II. Cells in D-E were from Fura-2 AM-stained slices and perfused with 4 mmol/l K^+^.

As observed for the higher concentration (Fig. 3D-G), TTA-P2 alone only incompletely reduced oscillatory activity (Fig. 5B-C; Supplementary Table S17-18; absolute data in Supplementary Fig. S5A and Table S19-20) and the number of active cells (Supplementary Fig. S5B) but increased the intra-burst frequency (Supplementary Fig. S5C-D; Supplementary Table S19 and 21). Addition of isradipine led to a cessation of all signals, demonstrating that the combination of both, T- and L-type VGCCs, is necessary for calcium oscillations and cannot be substituted for by other calcium channels.

This does not only extend to the oscillations of calcium but to calcium levels in general. Recording mean [Ca^2+^]_int_ levels using Fura-2 AM revealed that, upon inhibition of T- and L-type channels, calcium levels were undistinguishable from levels before stimulation with Ang II (Fig. 5D-E; Supplementary Table S22-23) but significantly lower than in controls (Supplementary Fig. S6A-B; Supplementary Table S24-25).

### L-type calcium channels are expressed in all ZG cells

The heterogeneous nature of the effects of TTA-P2 and isradipine led us to investigate whether differences in the expression of the main L- and T-type VGCCs exist across ZG cells. For this, we obtained access to a single-nuclear RNA-seq data set of mouse adrenal glands (unpublished data by AKS and UIS). We selected ZG cells based on the expression of *Dab2, Cacnb2, Agtr1b* and *Agtr1a* (see supplementary methods). A histogram of the normalized counts of transcripts for the three main VGCCs in the ZG (Cacna1c, Cacna1d and Cacna1h) revealed expression of Ca_V_1.2 in virtually all cells. About 20% of the cells, however, lacked expression of Cacna1d and/or Cacna1h (Fig. 6A). Plotting the normalized, natural-log transformed counts of transcripts for the T-type channel Ca_V_3.2/*Cacna1h* versus the L-type channel Ca_V_1.3/*Cacna1d* reveals four subsets of cells (Fig. 6B). The first consists of cells without detected RNA transcripts for either channel (red 1: 2252/6589 cells; 34%). There are also subsets of cells without detected *Cacna1d* but with *Cacna1h* transcripts (red 2: 1353 cells; 21%) and cells expressing *Cacna1d* without detected *Cacna1h* (red 3: 1729 cells; 26%). A similar number of cells expressed both, *Cacna1d* and *Cacna1h* (red 4: 1255 cells; 19%).

**Figure 6.**
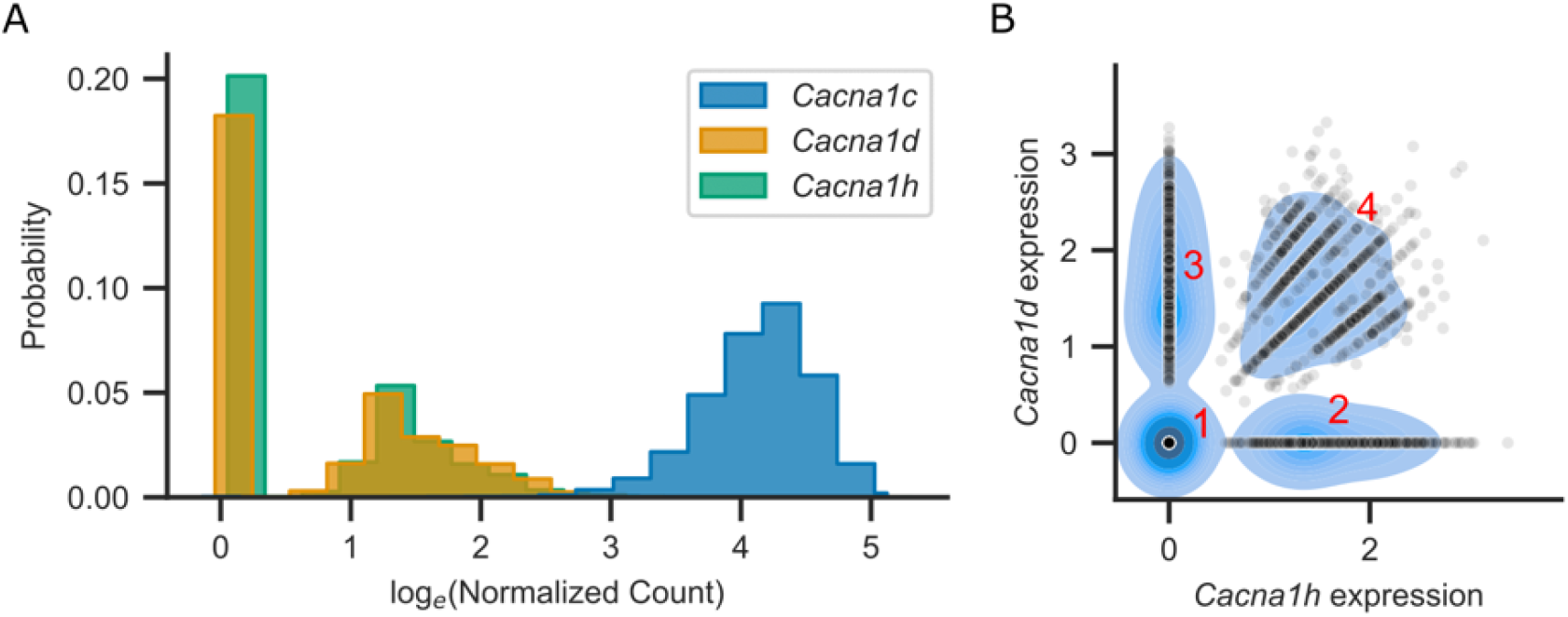
All ZG cells have transcripts for Ca_V_1.2 but not necessarily Ca_V_1.3 or Ca_V_3.2. **(A)** Histogram of the natural-log transformed normalized counts of indicated VGCC transcripts found in the ZG. Almost all cells exhibit expression of Cacna1c but not Cacna1d or Cacna1h. Bars are slightly shifted for each isoform to improve visibility. **(B)** Plot of the normalized, natural-log transformed counts of *Cacna1d* (Ca_V_1.3) versus *Cacna1h* (Ca_V_3.2) as individual points per cell. A kernel density plot of the data is plotted below in blue with darker color indicating higher density of values underneath. Red numbers indicate expression clusters explained in the text. The striped appearance in cluster 4 is due to the low value of *Cacna1h* and *Cacna1d* counts as seen in *A*.

## Discussion

The zona glomerulosa is a complex system with several stimuli regulating the synthesis of aldosterone^2^. It has been postulated that most of these factors, including serum potassium and Ang II, act by mediating oscillatory calcium influx into ZG cells elicited through depolarization of the cell membrane^2^ requiring Ca_V_3.2^3^ and potentially also through release of calcium from intracellular stores^26,27^. In this manuscript, we demonstrate that Ang II and potassium dependent [Ca^2+^]_int_ oscillations still occur in Ca_V_3.2 knock out mice and become instead dependent on L-type calcium channels. Furthermore, acute inhibition of T-type calcium channels does not abolish Ang II dependent calcium influx into the wild-type ZG. Rather, a subset of cells still exhibits oscillatory calcium influx via L-type calcium channels.

Hu et al. demonstrated that murine ZG cells constitute electrical oscillators with action potential-like depolarizations. Maintenance of these fluctuations were critically dependent on the voltage-gated T-type calcium channel Ca_V_3.2^3^. Although not proven so far, it is likely that these voltage oscillations are the basis of ZG calcium oscillations as they exhibit similar frequencies and stimulus dependence (Fig. 1)^7–9^. However, it is not the change in voltage but ultimately the influx of calcium that is required for the physiological function of the ZG. Contrary to this model, Ca_V_3.2 knock-out mice did not show reduced aldosterone levels^8,14^. However, these studies investigated KO mice on a systemic level, and secondary mechanisms (such as activation of the renin-angiotensin system or upregulation of other calcium influx pathways in the ZG) may have compensated for the loss. We therefore investigated the molecular mechanisms regulating calcium influx in the ZG of mice lacking the Ca_V_3.2 channel in comparison to WT mice in acute slice preparations and isolated adrenocortical cells.

We found that calcium signaling was largely unaffected by the loss of Ca_V_3.2 in KO mice (Fig. 1). Despite a slight increase in the spiking activity, mean levels of [Ca^2+^]_int_ in the ZG were indistinguishable from WT, explaining why aldosterone levels were unchanged (Fig. 1D and E). It was suggested that other T-type channels may also be expressed in the adrenal cortex^14^. However, our data from RNA-seq (Fig. 2A) did not suggest relevant expression of T-type channels other than *Cacna1h*. Adrenocortical cells from KO mice lacked T-type currents (Fig. 2D-E). Furthermore, perfusion of slices from adrenals of KO mice with the T-type inhibitor TTA-P2 did not result in changes to calcium signaling (Fig. 3), supporting that Ca_V_3.2 is indeed the only relevant T-type VGCC in the murine ZG.

On the other hand, perfusion with the L-type calcium channel inhibitor isradipine almost completely suppressed [Ca^2+^]_int_ oscillations in KO mice (Fig. 4), demonstrating that these signals instead depend on L-type calcium channels. Based on previous knowledge^3,19,20,28^ and our bulk RNA-seq analysis (Fig. 2A), the L-type calcium channels most prominently expressed in the murine and human adrenal cortex are Ca_V_1.3 and Ca_V_1.2^3,19,20^. We did not find evidence of compensatory upregulation of L-type or any other calcium channels or transporters in KO mice. Patch clamp experiments also ruled out large post-translational changes to voltage-gated calcium channels as current amplitudes and voltage-dependencies were similar in WT and KO mice (Fig. 2D-E).

The importance of L-type channels had already been previously demonstrated by studies identifying somatic gain-of-function Ca_V_1.3 mutations in patients with primary aldosteronism ^19,21,28–30^. Furthermore, there are previous reports that aldosterone synthesis in human ZG is dependent on both, T- and L-type, calcium channels^20,31^. It is currently not known whether human ZG cells exhibit similar voltage and [Ca^2+^]_int_ oscillations as murine ones. However, species differences in the ion channels involved seem to be mainly in the composition of potassium channels^32^, while the expression of calcium channels is rather similar^19,20,32,33^.

Currently, we cannot explain why [Ca^2+^]_int_ oscillations were faster in KO than in WT mice. It had already been suggested that additional conductances must underlie the initial depolarization in WT^3^, and this is confirmed by our results, as L-type channels (Ca_V_1.3 and even more so 1.2) require stronger depolarization for activation than T-type VGCCs and are unlikely to drive depolarization from a potassium-defined resting membrane potential on their own. Clearly, the closure of TASK potassium channels is involved in permitting (Ang II-dependent) depolarization^5,6,34^, but still other, currently unknown, conductances must mediate the initial depolarization itself.

Also, it is currently unclear how individual spikes and bursts are terminated. We previously observed slower spiking in a gain-of-function *Cacna1h* knock-in model. Given that the knock-out presented here should generally lead to the opposite effect, it is a tempting speculation that the frequency of calcium spiking is inversely linked to intracellular [Ca^2+^]. In our knock-out model this would serve to sustain physiological calcium levels (and hence aldosterone synthesis). Previous observations have suggested a role for calcium-dependent potassium channels^35,36^ in the ZG, which may control spiking, but further work is necessary to understand their functional importance. This would best combine *in situ* electrophysiological studies with simultaneous calcium imaging, which we were unfortunately unable to perform here.

Besides the situation of a chronic loss of Ca_V_3.2 as in our KO model, our data also imply that physiological calcium signaling in the WT ZG is not solely dependent on Ca_V_3.2. Acute inhibition of T-type VGCCs with the specific blocker TTA-P2 in WT mice only attenuated activity by approximately 50-80 % (Fig. 3&5). [Ca^2+^]_int_ oscillations could only be completely inhibited when simultaneously blocking L-type calcium channels using isradipine (Fig. 5). Acute inhibition of L-type channels in isolation, however, did not alter calcium spiking (Fig. 4). This suggests that the generation of oscillations in WT ZG cells is primarily dependent on Ca_V_3.2. However, following the acute inhibition of T-type calcium channels, for example via TTA-P2, L-type channels can maintain some degree of calcium signaling in WT cells (Fig. 3). Cells with remaining activity after inhibition of Ca_V_3.2 exhibited higher intra-burst frequencies than before inhibition (Fig. 3G; Supplementary Fig. S3B), suggesting that a similar mechanism as in KO cells is also present in this subset of WT cells.

Our analysis of murine single-nuclear RNA-seq data suggests that Ca_V_1.2 is expressed in all ZG cells while transcripts for Ca_V_1.3 were not found in all ZG cells (Fig. 6). The latter could be due to lower expression levels and low sensitivity of single-nuclear sequencing. It remains puzzling why no L-type currents were observed in ZG cells in the study by Hu et al., even when stimulated with the L-type channel activator Bay K8644^3^. In contrast, we observed L-type currents in all cells recorded using whole cell patch-clamp, even without further stimulation (Fig. 2E). While this discrepancy might be explained by the difference in cell preparation (in-situ patch clamp^3^ versus dissociated adrenocortical cells in our study), our calcium imaging experiments and results from single-nuclear RNA-seq (Fig. 6) also clearly support the relevance of L-type calcium channels in generating and maintaining calcium oscillations in the ZG of both genotypes, WT and KO, *in situ*.

Furthermore, we could demonstrate that Ang II mostly changes [Ca^2+^]_int_ through variations of the patterns of oscillations and not by altering baseline levels (Fig. 5D-E). Simultaneous inhibition of T- and L-type calcium channels not only completely stopped Ang II dependent [Ca^2+^]_int_ oscillations but also any changes in baseline [Ca^2+^]_int_. This suggests that intracellular calcium stores may not play a large role in Ang II-dependent aldosterone synthesis but further studies are required.

Besides unraveling the physiology of calcium signaling in the ZG leading to aldosterone productions, our findings also have implications in directing future pharmacological interventions. It has been suggested that the inhibition of T-type calcium channels might represent a promising target for treating hyperaldosteronism (or lowering aldosterone synthesis) in general^13^. Our results, however, imply that a long-term treatment would likely be countered via calcium influx through L-type calcium channels. This explains why mibefradil, a preferential T-type channel inhibitor, did not exert persistent effects on aldosterone levels or blood pressure *in vivo*^37,38^. Nevertheless, a specific blockade of Ca_V_3.2 or Ca_V_1.3 in primary aldosteronism due to gain-of-function mutations in either channel may still constitute a valid therapeutic strategy to alleviate pathological calcium influx into the ZG^8,19,28^. Interestingly, it has been observed in the H295R cell line that a blockade of both, L- and T-type channels (for example using the unspecific inhibitors benidipine^39^ or efonidipine^40^) was efficient in reducing aldosterone production. This may indeed be a strategy to lower aldosterone synthesis, but potential extra-adrenal side effects may limit their usefulness, and this approach therefore requires further investigation.

## Acknowledgements

We would like to thank the team of the Core Unit Genomics of the Berlin Institute of Health at Charité – Universitätsmedizin Berlin, in particular Dr. Tatiana Borodina, for performing library preparation and sequencing of our RNA samples. We are very grateful to Sarah Döring, Nico Brüssow, Marie Cotta and Ana Lucia Huitron Carrizales for performing mouse genotyping. The authors also would like to thank Drs. Julia Schewe and Eric Seidel for their help with establishing the Cacna1h^-/-^ colony.

## Author Contributions

G.S. conceived the study; H.A.D. and M.V. performed calcium imaging; H.A.D. and G.S. analyzed calcium imaging data; H.A.D. and G.S. analyzed bulk RNA-seq data; H.A.D. performed and analyzed patch clamp experiments of adrenocortical cells; A.K.S. and U.I.S. designed single nucleus RNA-seq experiments; A.K.S. performed and analyzed single nucleus RNA-seq experiments; H.A.D, U.I.S. and G.S. wrote the manuscript with contributions from all authors.

