## Supplementary Information for "T- and L-type calcium channels maintain calcium oscillations in the murine zona glomerulosa"

#### Detailed Methods

##### Generation and breeding of the *Cacna1h* KO mouse model

*Cacna1h* KO mice (based on C57BL/6J strain) carrying an 8-bp deletion on exon 25, with subsequent frameshift and termination (p.His1570Glnfs*83, *Cacna1h*$^{+/-}$) were generated previously[1]. *Cacna1h* KO and control mice were bred and housed at the Forschungseinrichtungen für Experimentelle Medizin (FEM; Charité - Universitätsmedizin Berlin) under specific-pathogen free conditions in a 12-h light/dark cycle with *ad libitum* access to food and water.

C57BL/6J mice for single cell RNA-seq experiments were bred and housed at Max Planck Institute for Molecular Genetics (Berlin) and maintained under specific pathogen-free conditions in a 12h light/dark cycle with *ad libitum* access to food and water.

All animal experiments were approved by the local authority (Landesamt für Gesundheit und Soziales Berlin) and performed under consideration of all relevant ethical regulations.

##### Organ harvest

To harvest adrenal glands for experiments, 10 to 30 week-old mice were anesthetized using isoflurane (400 µL as open drop in a 2-l beaker) and euthanized by cervical dislocation. The abdominal cavity was then opened using scissors, and the adrenal glands identified and rapidly extracted. The procedure of organ harvest following euthanasia was performed according to German law and registered with the local authorities (Landesamt für Gesundheit und Soziales (LaGeSo) Berlin; T 0425/17).

##### Calcium imaging

For acute slice preparations, similar to previously published procedures[1,2], adrenal glands were transferred into ice cold bicarbonate-buffered saline (BBS: 100 mmol/l NaCl, 2 mmol/l KCl, 26 mmol/l $NaHCO_3$, 0.1 mmol/l $CaCl_2$, 5 mmol/l $MgCl_2$, 10 mmol/l D-glucose, 10 mmol/l HEPES) immediately after extraction. During all following steps, BBS was continuously gassed with carbogen (95 % $O_2$ + 5 % $CO_2$). Both glands were embedded into 4 % low-melting agarose, following the removal of the surrounding fat. The organs in the agarose block were glued to the mounting stage of a vibratome (7000 smz-2, Campden Instruments) and slices were cut at 120 µm thickness. During cutting, adrenal glands in the agarose block and slices were maintained in ice-cold, gassed BBS. Slices were transferred into BBS at 35°C for 15 minutes for regeneration and subsequently stored for up to 6h at RT in BBS supplemented with 2 mmol/l $CaCl_2$.

Slice staining was performed in a cell culture insert within a single well of a 24-well plate at room temperature. The insert was filled with 250 µL containing initially 37 µmol/l Calbryte 520 AM and

0.001 % Pluronic F-127 or 64 µmol/l Fura-2 AM and 0.16 % Pluronic dissolved in BBS (supplemented with 2 mmol/l $CaCl_2$). The well was filled with 750 µL BBS (supplemented with 2 mmol/l $CaCl_2$) and continuously gassed with carbogen. The slices were incubated in the staining solution for 1 h. Following Fura-2 AM staining, slices were transferred to fresh BBS (without Fura-2, supplemented with 2 mmol/l CaCl2) for 15 min de-esterification.

For recording, slices were placed in a recording chamber and were continuously perfused with solution from a reservoir. The solution reservoir was supplied with carbogen gas and was heated via an inline heating coil to a temperature of 30 ± 1°C. Two solutions were prepared and mixed in order to yield the potassium concentration of interest (BBS $2K^+$ : 100 mmol/l NaCl, 2 mmol/l KCl, 26 mmol/l $NaHCO_3$, 5 mmol/l NaGluconate, 2 mmol/l $CaCl_2$, 1 mmol/l $MgCl_2$, 10 mmol/l D-glucose, 10 mmol/l HEPES); and BBS $7K^+$: 100 mmol/l NaCl, 2 mmol/l KCl, 26 mmol/l $NaHCO_3$, 5 mmol/l KGluconate, 2 mmol/l $CaCl_2$, 1 mmol/l $MgCl_2$, 10 mmol/l D-glucose, 10 mmol/l HEPES)). Ang II was added from a 1 µmol/l stock to yield final concentrations as indicated.

Calbryte fluorescence was excited at 470 nm while Fura-2 was alternately excited at 340 and 385 nm using a FuraLED light source (Cairn Research). Images were taken every 100 ms with 10-ms exposure using an OptiMOS camera (QImaging).

**Electrophysiological recordings of dissociated adrenal cortical cells**

For dissociated adrenal cortical cell preparations, extracted adrenal glands were rapidly transferred into bath solution containing 140 mmol/l NaCl, 4 mmol/l KCl, 1 mmol/l $MgCl_2$, 2 mmol/l $CaCl_2$, 10 mmol/l HEPES on ice. After the removal of the surrounding fat, the glands were cut in half and the medulla manually removed. The glands were further cut into fragments (< 0.5 mm) and incubated in bath solution supplemented with collagenase type I (1mg/ml), collagenase type IV (1mg/ml), penicillin (50 U/ml) and streptomycin (50 mg/ml) for 20 min at 37°C on a shaker set to 300 rpm. Subsequently, shear-based dissociation of the cells was performed in a 1 ml syringe attached to a cannula (18-20 G) and polyethylene tubing (30 cm) of matching diameter. A syringe pump was used to provide a controlled flow rate (700 µL/min) of the cell suspension through the cannula and tubing. The cell suspension was centrifuged for 5 min at 300 g and room temperature. The cell pellet was resuspended in DMEM/F12 (1:1) supplemented with Ultroser G (2 %), penicillin (50 U/ml) and streptomycin (50 mg/ml), α-tocopherol (1 µmol/l), insulin transferrin selenium (1 %), ascorbic acid (100 µmol/l). Cells were seeded on coverslips (9 mm diameter, coated with laminin and poly-L-lysin) in a 24-well plate, incubated in a 5 % $CO_2$ humidified atmosphere at 37°C and used for whole-cell patch clamp recordings the next day.

Currents were recorded on a HEKA EPC 10 amplifier (HEKA Elektronik) using Patchmaster software. Borosilicate pipettes with resistances of 2–4 MΩ were pulled on a Sutter P-1000 puller (Harvard

Apparatus) and subsequently fire polished using a Narishige MF-890 microforge. Extracellular solution contained 100 mmol/l NaCl, TEACl 20 mmol/l, 4 mmol/l KCl, 1 mmol/l $MgCl_2$, 10 mmol/l HEPES, 10 mmol/l $CaCl_2$, pH 7.4. Pipette solution contained 140 mmol/l CsCl, 10 mmol/l NaCl, 5 mmol/l EGTA, 2 mmol/l $MgCl_2$, 10 mmol/l HEPES, pH 7.4. Currents were elicited by voltage steps ranging from either -95 or -55 to 75 mV in 10 mV increments from a holding potential of -80 mV or -40 mV, respectively.

The obtained data was fit with equation 1:

$$[1] \qquad I(V) = G * (V - V_{rev}) * \frac{1}{1+e^{\frac{-(V-V_{0.5})}{k}}}$$

where *I* is the macroscopic current, *G* the maximum conductance, *V* the tested voltage, $V_{rev}$ the reversal potential, $V_{0.5}$ the half-maximal activation and k the slope.

**Bulk RNA-seq**

For RNA extraction from whole adrenal cortices, extracted adrenal glands from 6 WT and 6 KO (3 male, 3 females each) were rapidly transferred into PBS on ice. After the removal of the surrounding fat, the glands were cut in half, manually separated from the medulla and placed in RNAlater (ThermoFisher). Total RNA was isolated using the RNeasy Mini Kit (Qiagen) with subsequent DNAse digestion according to the manufacturer's instructions. Purified RNA was then further processed at the Core Unit Genomics of the Berlin Institute of Health at Charité – Universitätsmedizin Berlin. Ribosomal RNA was depleted and cDNA transcribed using the TruSeq Stranded Total RNA kit (Illumina). 150bp paired-end Sequencing was performed on a NovaSeq 6000 device on a single lane of a SP flow cell (all from Illumina).

The complete bioinformatics pipeline used in this study, including all parameters used, can be found on GitHub (https://github.com/dhoangan/Cacna1hKO). In brief, results were analyzed on the background of Ensembl release 106 of the GRCm39 cDNA transcriptome. Indexing of the transcriptome was performed using Salmon[3] (version 1.5.2). Initial quality control of the results was performed using MultiQC[4] (version 1.11). Further quality checks, filtering and trimming was performed using fastp[5] (version 0.12.4). Salmon was also used for mapping of reads. Following inspection, one male KO sample had to be excluded from further analysis as ribosomal RNA made up more than 80 % of all reads, likely because rRNA depletion in the TruSeq Kit failed.

Raw data from our bulk RNA-seq experiment can be accessed at ArrayExpress with the accession number E-MTAB-12999 (https://www.ebi.ac.uk/biostudies/arrayexpress/studies/E-MTAB-12999).

Filtered gene counts were further processed and analyzed using DESeq2[6] (version 1.38.3) in R (version 4.2.1). Differentially expressed genes (DEGs) were identified using the DESeq function with the chosen threshold criteria of p-values < 0.05 (adjusted using the Benjamini–Hochberg correction) and a 2-fold change (log2 transformed fold change ± 1, after shrinkage using the apeglm method[7]).

For heatmap generation, raw read counts were normalized using the DESeq2. If indicated, a Z-score normalization was performed on the normalized read counts across samples for each gene.

Functional enrichment analysis of the identified DEGs was assessed based on Gene Ontology[8] (GO) and KEGG pathway[9] over-representation using the ClusterProfiler[10] package (version 4.6.2) in R. A p-value less than 0.05 was considered as statistically significant.

**Single cell RNA-seq**

After adult mice (two males and two females; each seven weeks old) were killed by cervical dislocation, both adrenal glands were collected, snap frozen in liquid nitrogen and stored at -80°C until further procedures.

Adrenal glands were processed using a dounce homogenizer in a Triton X-100 lysis buffer (10% Triton X-100, Superasin 20 U/µL, RNAseIn U/µl, Nuclei Isolation Medium: 1mM DTT, 50X Protease Inhibitor, 1.5M Sucrose, 1M KCL, 1M $MgCl_2$, 1M Tris buffer pH 8.0) to get a nuclei suspension. Subsequently, the suspension was stained with DAPI (4',6-diamidino-2-phenylindole) and sorted by flow cytometry (BD FACSAria III High Sensitivity Flow Cytometer (mouse), BD Genomics, USA) into RNAse inhibitor containing tubes. The nuclei quality was assessed under Leica DMi8 microscope and quantified using a Neubauer chamber.

Each single-nuclei suspension was loaded onto a Chromium Single Cell 3'Chip A (10X Genomics) to yield GEMs (Gel Beads in Emulsion). Subsequently, GEMs were processed according to Chromium Single Cell 3' Reagent Kits v3 (CG000183 Rev C), to generate sequencing-ready libraries. Libraries were sequenced on an Illumina HiSeq 4000 following 10X Genomics sequencing guidelines.

All count matrices were processed with Seurat using *CreateSeuratObject* function (*min.cells = 3, min.features = 200*): further filtering was performed during QC step, by filtering cells less than 700 genes (*nFeature_RNA > 700*) and outliers with over 10,000 UMIs were removed (*nCount_RNA < 10,000*). Next, filtered matrices were merged (ncells = 27,448) and normalization was performed using *SCTransform*(2), regressing out cell cycle and the overexpressed *Malat1* gene (*vars.to.regress = c("malat","S.Score","G2M.Score"*) in the process. After PCs were computed (*RunPCA*), UMAP was generated with the first 30 PCs (*RunUMAP*). Clustering and cell type identification was conducted using cNMF(3): briefly, a total of 10 gene expression programs (GEPs) were produced, and a "usage" matrix was created to indicate the extent to which each GEP contributes to a cell's identity. Each cell was assigned a discrete cluster identity based on the highest usage score obtained. Differentially expressed genes were then computed using *FindAllMarkers* (*test.use = "LM"*, *latent.vars = "nCount_RNA"*), and the annotations of each cluster was done by cross-referencing their top expressed genes with established markers from literature.

Gene expression program 8 had canonic ZG markers such as *Dab2* (Disabled homolog 2), *Cacnb2* (Calcium voltage-gated channel auxiliary subunit beta 2), *Agtr1b* (Angiotensin II receptor, type 1b) and *Agtr1a* (Angiotensin II receptor, type 1b), therefore we identified cells with a high GEP 8 score as ZG. Using the subset() function, we extracted 6589 ZG cells.

**Statistics**

All P-values (except for the differential expression analysis) are results from a likelihood ratio test of the linear mixed models and are indicated as follows: ns, $P \geq 0.05$; *, <0.05; **, P<0.01; ***, P<0.001. Statistical mixed model analysis was performed in R (4.2.1) using the glmmTMB package (version 1.1.5). Normality of the data was assured visually using quartile-quartile plots and inspection of residuals after fitting. If data was not normally distributed, a diverse set of distributions was tested using DHARMa (version 0.4.6), and either the log-Tweedie or log-Gamma distribution was chosen for further analysis using glmmTMB. Individual recordings were treated as random effect. All error bars and bands show 95% confidence interval of the mean value.

For differential expression analysis, reads obtained by RNA sequencing were normalized and differentially analyzed using the DESeq2 package (version 1.38.3). Genes were considered as significantly regulated with the chosen threshold criteria of a 2-fold change (log2-transformed fold change ± 1) and a P-value lower than 0.05 (adjusted by Benjamini–Hochberg procedure for multiple comparisons).

If not otherwise mentioned, larger and smaller spots in figures represent mean individual values per slice and cell, respectively. The mean of all mean values per slice are shown as superimposed large white circles ± 95% confidence interval (error bars, CI). Shades areas and superimposed larger circles represent error bands indicating ± 95% CI and mean values of cells, respectively (Fig. 1D, 1E, 3E, 4C, 5B, 5D). Confidence intervals were calculated from a bootstrap resampling.

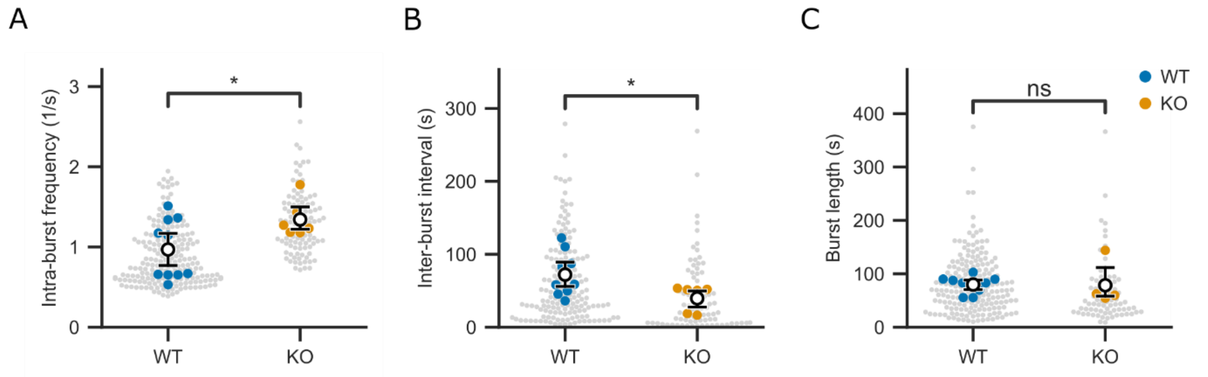

***Supplementary Figure S1.*** *Calcium bursting analysis of data obtained from adrenal slices of WT (blue) or KO (orange) mice stained with Calbryte 520 AM in the presence of 4 mmol/l K+ and 500 pmol/l Ang II.* ***(A)*** *Average frequencies of calcium oscillations during bursts (intra-burst frequency) were higher in KO ZG cells than WT.* ***(B)*** *The average time not spent in calcium bursting (inter-burst interval) was lower in KO cells than WT.* ***(C)*** *The average length of a burst was unchanged between genotypes. P values (likelihood ratio test of linear mixed models) are indicated as follows: ns, P ≥ 0.05; *P < 0.05; See Supplementary Table S1 for statistical information. All data are shown as mean (white circle) ± 95 % confidence intervals (CI, error bars).*

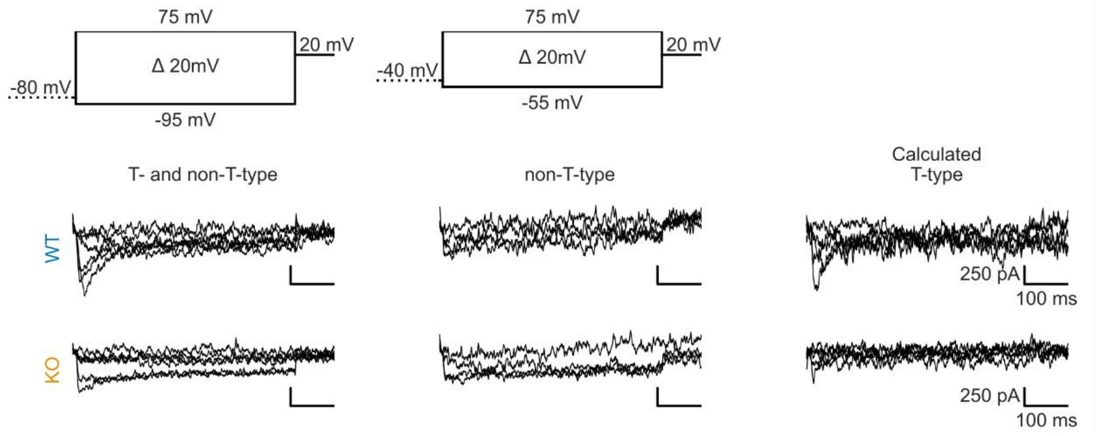

***Supplementary Figure S2.*** *Representative recordings of the calcium currents in isolated adrenocortical cells from WT and KO mice elicited by two voltage clamp protocols: (i) starting from a holding potential at - 80 mV with voltage steps between -95 and +75 mV to evoke calcium currents (upper, left), (ii) from a holding potential at - 40 mV with voltage steps between -55 and +75 mV (upper, middle). The resulting T-type current (right) was calculated by subtracting the current elicited from -40 mV (middle) from the one from -80 mV (left).*

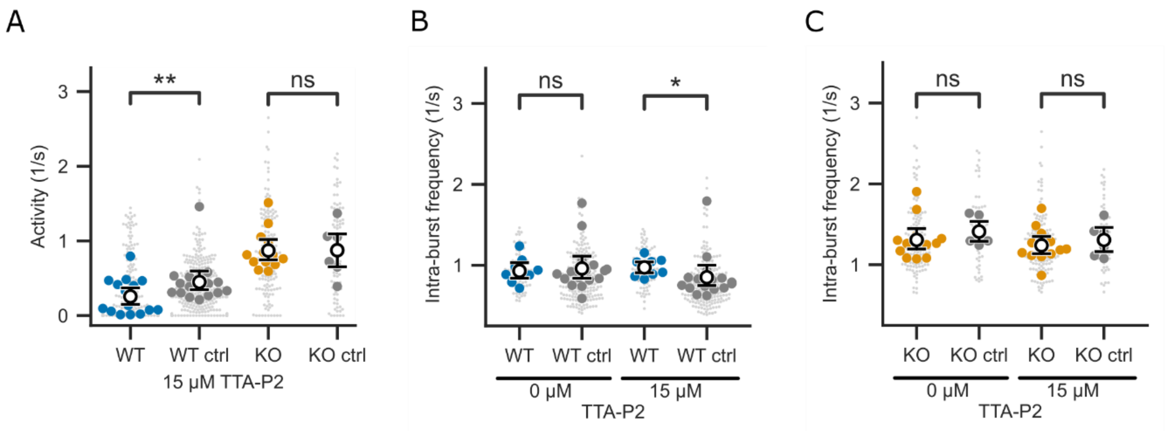

***Supplementary Figure S3. (A)*** *Calcium spike activity was significantly lowered by 15 µmol/l TTA-P2 in WT (blue) but not KO (orange) compared to corresponding control.* ***(B, C)*** *WT but not KO cells showed a significantly changed intra-burst frequency upon perfusion with 15 µmol/l TTA-P2, when compared to corresponding controls. Only cells that remained active at 15 µmol/l TTA-P2 are shown in B and C. Data in A-C are shown as overall mean per group (white circle) ± 95% CI (error bars). Individual values of cells and slices are shown as smaller and larger spots, respectively. Values were determined at time span -2 - 0 and 5.5 - 7.5 minutes before and after addition of inhibitor, respectively (indicated by the black dashed and solid line in 3E). P values (likelihood ratio test of linear mixed models) are indicated as follows: ns, P ≥ 0.05; *P < 0.05; **P < 0.01; ***P < 0.001. See Supplementary Table S10-12 for statistical information.*

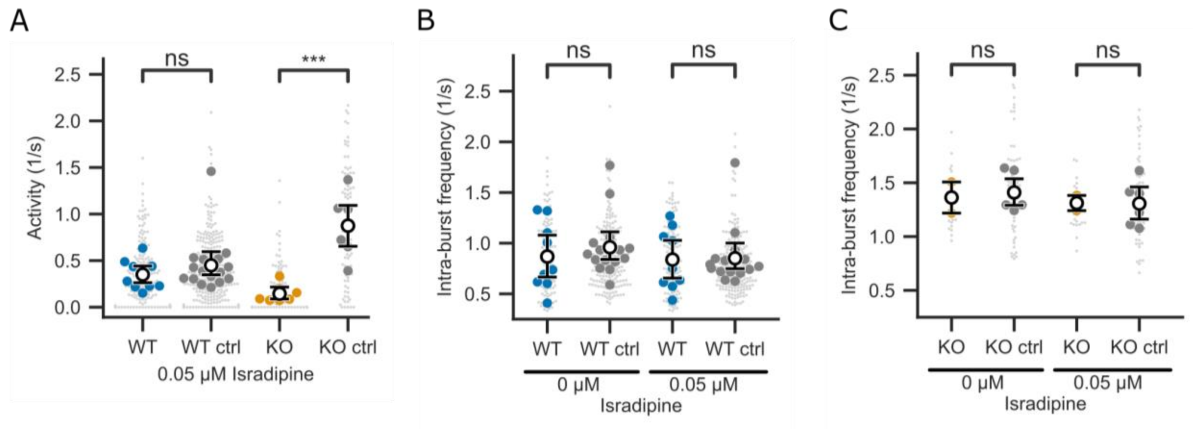

***Supplementary Figure S4. (A)*** *Calcium spike activity was significantly lowered in KO but not WT by 0.05 µmol/l isradipine.* ***(B-C)*** *The intra-burst frequency was unchanged upon perfusion with 0.05 µmol/l isradipine in WT and KO cells compared to control. Only cells that remain active at 0.05 µmol/l isradipine are shown in B and C. Data in A-C are shown as overall mean per group (white circle) ± 95% CI (error bars). Individual values of cells and slices are shown as smaller and larger spots, respectively. P values (likelihood ratio test of linear mixed models) are indicated as follows: ns; ***P < 0.001. Values were determined at time span -2 - 0 and 5.5 - 7.5 minutes before and after addition of inhibitor, respectively (indicated by the black dashed and solid line in 4C). See Supplementary Table S14-16 for statistical information.*

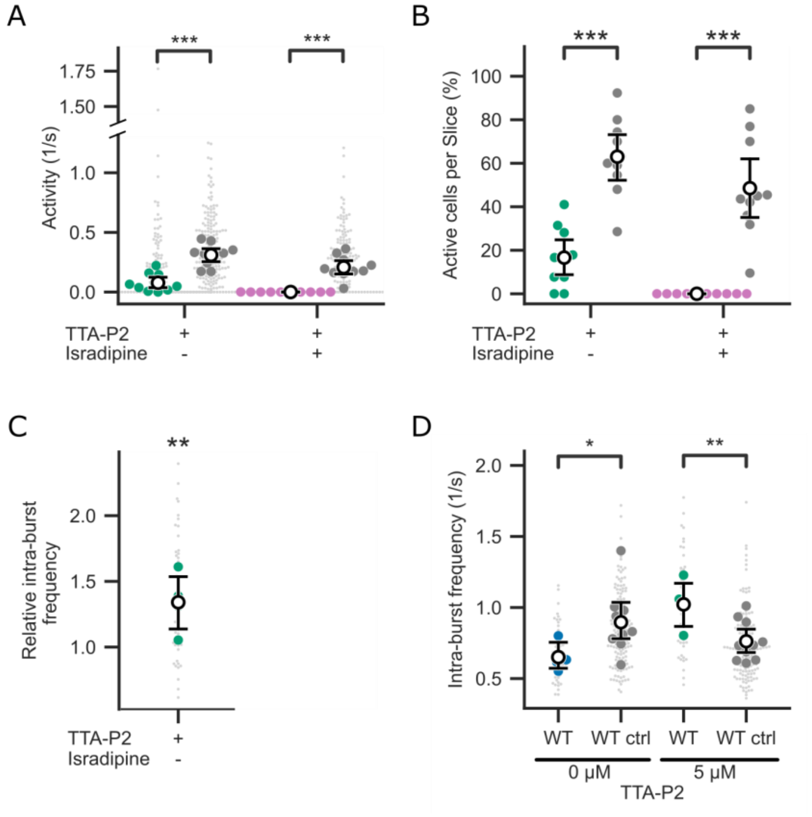

***Supplementary Figure S5. (A)*** *Calcium spike activity in cells from WT adrenal slices stained with Calbryte 520 AM at 4 mmol/l K*$^{+}$ *and 500 pmol/l Ang II was lowered by 5 µmol/l TTA-P2 alone (green) and in combination with 300 nmol/l isradipine (pink) compared to controls (grey, no inhibitors).* ***(B)*** *Fraction of cells in a WT adrenal slice that were active during the steady-state phases as indicated in 5B compared to untreated controls (grey).* ***(C)*** *Intra-burst frequency of WT cells during the green steady-state phase as indicated in 5B relative to controls.* ***(D)*** *Intra-burst frequency of the remaining active WT and WT control cells at 0 and 5 µmol/l TTA-P2. Individual larger circles show mean values of slices (green or pink: WT; grey: WT control). Smaller spots in A, C and D show individual mean values of cells. White circles show overall mean per group of mean values of slices (errorbars: ± 95% CI). For control, cells were perfused without calcium channel inhibitors. The color of the mean values correspond to the color-coded intervals in Fig. 5B. P values (likelihood ratio test of linear mixed models) are indicated as follows: ns, P ≥ 0.05; *P < 0.05; **P < 0.01; ***P < 0.001. See Supplementary Table S18-21 for statistical information.*

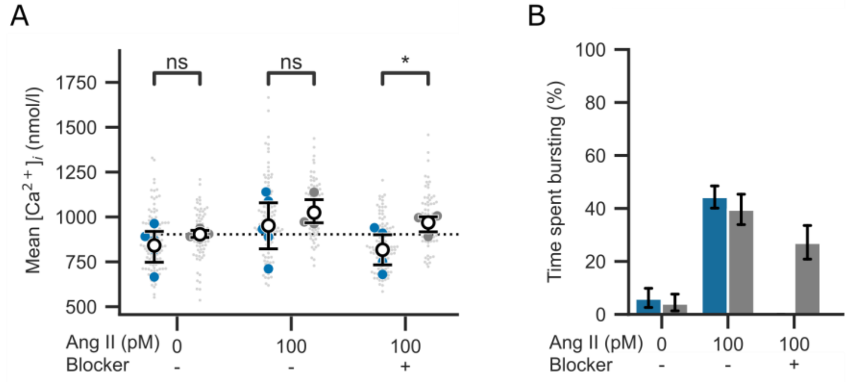

***Supplementary Figure S6. (A)*** *Mean intracellular calcium concentrations of ZG cells from Fura-2-AM-stained adrenal slices before and during perfusion with both inhibitors TTA-P2 (5 µmol/l) and isradipine (300 nmol/l) at varying concentrations of Ang II as indicated in 5D (black bars; blue circles: treated WT; grey circles: untreated*

*WT controls). Mean values for each slice and cell are shown as larger and smaller circles, respectively. Overall means per group are shown as white circles ± 95% CI.* ***(B)*** *Time spent bursting of ZG cells from Fura-2-AM-stained adrenal slices before and during perfusion with both inhibitors TTA-P2 (5 µmol/l) and isradipine (300 nmol/l) at varying concentrations of Ang II as indicated in 5E (blue bars: treated cells, grey bars: untreated controls, error bars: 95% CI). See Supplementary Table S24-25 for statistical information.*

**Table S1. Calcium bursting parameters (Fig. 1 and S1)**

| | WT | | | KO | | | |
|---|---|---|---|---|---|---|---|
| | Mean | CI (lower limit) | CI (upper limit) | Mean | CI (lower limit) | CI (upper limit) | p-value |
| Activity (1/s) | 0.494 | 0.368 | 0.631 | 1.054 | 0.898 | 1.220 | 0.003, ** |
| Intra-burst frequency (1/s) | 0.969 | 0.777 | 1.173 | 1.343 | 1.223 | 1.500 | 0.013, * |
| Inter-burst interval (s) | 73.823 | 58.706 | 89.901 | 39.596 | 27.559 | 49.615 | 0.013, * |
| Burst length (s) | 80.718 | 72.233 | 88.582 | 69.630 | 51.174 | 96.538 | 0.242, ns |
| n(animals) | 7 | | | 5 | | | |
| n(slices) | 11 | | | 7 | | | |
| n(cells) | 208 | | | 103 | | | |

***Supplementary Table S2. Mean calcium concentration at 100 pmol/l Ang II and indicated $K^+$ concentrations in nmol/l (Fig. 1D).***

| | WT | | | | | | KO | | | | | | |
|---|---|---|---|---|---|---|---|---|---|---|---|---|---|
| $K^+$ (mmol/l) | Animals | Slices | Cells | Mean | 95% CI (lower limit) | 95% CI (upper limit) | Animals | Slices | Cells | Mean | 95% CI (lower limit) | 95% CI (upper limit) | p-value |
| 3.0 | 3 | 4 | 57 | 673.109 | 640.176 | 705.589 | 7 | 11 | 162 | 645.48 | 680.019 | 748.391 | 0.571, ns |
| 3.5 | 4 | 6 | 97 | 769.015 | 741.215 | 796.764 | 6 | 9 | 85 | 714.374 | 782.856 | 858.993 | 0.367, ns |
| 4.0 | 4 | 6 | 89 | 826.939 | 802.065 | 852.085 | 7 | 12 | 102 | 821.389 | 740.414 | 829.582 | 0.848, ns |
| 4.5 | 4 | 6 | 120 | 835.522 | 805.055 | 866.717 | 5 | 6 | 73 | 784.304 | 868.558 | 936.949 | 0.521, ns |
| 5.0 | 4 | 6 | 81 | 937.409 | 893.847 | 982.227 | 5 | 6 | 91 | 902.708 | 839.545 | 903.299 | 0.374, ns |
| 5.5 | 3 | 6 | 52 | 930.973 | 871.769 | 990.086 | 6 | 9 | 99 | 870.952 | 863.787 | 991.088 | 0.556, ns |
| 6.0 | 3 | 6 | 88 | 1049.245 | 997.559 | 1103.995 | 3 | 5 | 59 | 924.61 | 680.019 | 748.391 | 0.316, ns |

***Supplementary Table S3. Mean calcium concentration at 4 mmol/l $K^+$ and indicated Ang II concentrations in nmol/l (Fig. 1E).***

| | WT | | | | | | KO | | | | | | |
|---|---|---|---|---|---|---|---|---|---|---|---|---|---|
| Ang II (pmol/l) | Animals | Slices | Cells | Mean | 95% CI (lower limit) | 95% CI (upper limit) | Animals | Slices | Cells | Mean | 95% CI (lower limit) | 95% CI (upper limit) | p-value |
| 20 | 8 | 7 | 116 | 744.666 | 714.61 | 778.21 | 8 | 9 | 96 | 720.323 | 689.729 | 750.844 | 0.259, ns |
| 50 | 7 | 10 | 127 | 737.938 | 708.891 | 768.572 | 5 | 5 | 53 | 678.321 | 629.572 | 728.491 | 0.766, ns |
| 70 | 7 | 13 | 173 | 812.12 | 787.717 | 836.24 | 8 | 12 | 94 | 815.891 | 783.031 | 850.125 | 0.749, ns |
| 100 | 5 | 12 | 85 | 871.254 | 839.655 | 904.306 | 9 | 16 | 119 | 807.653 | 774.643 | 841.156 | 0.363, ns |
| 200 | 9 | 6 | 131 | 880.731 | 848.105 | 913.548 | 11 | 16 | 172 | 841.667 | 810.604 | 872.044 | 0.296, ns |
| 300 | 8 | 14 | 103 | 903.257 | 874.88 | 932.768 | 9 | 13 | 140 | 867.233 | 835.629 | 900.215 | 0.175, ns |
| 500 | 7 | 8 | 155 | 867.103 | 836.186 | 898.888 | 10 | 14 | 159 | 821.247 | 789.112 | 855.352 | 0.496, ns |

**Supplementary Table S4. Normalized $log_2$ transformed read counts, the $log_2$ transformed fold-change and P-values of voltage-gated calcium channel transcripts detected in WT and KO samples (Fig. 2A)**

| | WT | | | | | | | KO | | | | | | Differential expression | | |
|---|---|---|---|---|---|---|---|---|---|---|---|---|---|---|---|---|
| | **S1** | **S2** | **S3** | **S4** | **S5** | **S6** | **mean** | **S7** | **S8** | **S9** | **S10** | **S11** | **mean** | **$Log_2$ fold-change** | **P-value** | **$P_{adj}$-value** |
| ***Cacna1a*** | 5.43 | 6.85 | 5.52 | 5.14 | 6.85 | 6.37 | 6.03 | 5.12 | 4.97 | 6.59 | 5.43 | 5.38 | 5.50 | -0.01426 | 0.164 | 1.000 |
| ***Cacna1b*** | 5.45 | 4.33 | 6.94 | 5.30 | 6.23 | 6.31 | 5.76 | 6.32 | 5.88 | 6.15 | 5.63 | 6.15 | 6.03 | 0.001699 | 0.876 | 1.000 |
| ***Cacna1c*** | 12.02 | 11.83 | 11.73 | 11.54 | 11.96 | 11.87 | 11.83 | 11.79 | 12.05 | 11.87 | 11.94 | 12.04 | 11.94 | 0.037301 | 0.268 | 1.000 |
| ***Cacna1d*** | 7.66 | 7.46 | 8.84 | 7.91 | 9.18 | 8.81 | 8.31 | 8.81 | 8.46 | 8.79 | 8.35 | 8.08 | 8.50 | 0.003163 | 0.818 | 1.000 |
| ***Cacna1e*** | 3.14 | 1.76 | 2.89 | 3.32 | 4.75 | 3.45 | 3.22 | 3.73 | 2.50 | 3.11 | 2.75 | 4.40 | 3.30 | -0.00007.2 | 0.991 | 1.000 |
| ***Cacna1f*** | 0.76 | 3.59 | -0.87 | 0.00 | 0.12 | 1.03 | 0.77 | 2.43 | 1.27 | 0.00 | 1.04 | -0.03 | 0.94 | -0.00074 | 0.835 | 1.000 |
| ***Cacna1g*** | 5.15 | 4.44 | 4.59 | 3.94 | 4.12 | 4.07 | 4.38 | 6.41 | 4.33 | 3.85 | 4.30 | 3.72 | 4.52 | 0.005836 | 0.492 | 1.000 |
| ***Cacna1h*** | 11.62 | 11.53 | 12.23 | 11.48 | 10.86 | 11.95 | 11.61 | 10.84 | 9.84 | 10.43 | 9.64 | 10.08 | 10.16 | -1.34825 | 0.000 | *** 0.000 |
| ***Cacna1i*** | 3.50 | 4.96 | 4.61 | 4.80 | 4.87 | 3.95 | 4.45 | 5.30 | 5.71 | 4.14 | 4.22 | 3.58 | 4.59 | 0.005232 | 0.593 | 1.000 |
| ***Cacna2d1*** | 12.84 | 12.40 | 12.79 | 12.59 | 13.19 | 12.85 | 12.78 | 13.22 | 13.08 | 13.24 | 13.05 | 13.16 | 13.15 | 0.239179 | 0.005 | 0.079 |
| ***Cacna2d2*** | 5.89 | 5.75 | 6.53 | 5.50 | 6.33 | 5.51 | 5.92 | 5.79 | 6.89 | 6.01 | 5.73 | 5.80 | 6.04 | 0.006066 | 0.644 | 1.000 |
| ***Cacnb1*** | 6.22 | 6.35 | 6.75 | 6.13 | 6.20 | 6.43 | 6.35 | 6.15 | 5.62 | 6.71 | 6.17 | 6.18 | 6.17 | -0.01216 | 0.420 | 1.000 |
| ***Cacnb2*** | 10.55 | 10.71 | 10.48 | 10.65 | 10.48 | 10.41 | 10.55 | 10.18 | 10.37 | 9.99 | 10.35 | 10.54 | 10.28 | -0.13882 | 0.018 | 0.263 |
| ***Cacnb3*** | 8.03 | 7.79 | 7.98 | 7.97 | 7.68 | 7.70 | 7.86 | 7.97 | 7.69 | 7.72 | 7.47 | 7.61 | 7.69 | -0.04515 | 0.159 | 1.000 |
| ***Cacnb4*** | 1.87 | 2.29 | 5.99 | 3.60 | 3.75 | 5.30 | 3.80 | 4.69 | 2.75 | 3.26 | 3.36 | 2.63 | 3.34 | -0.00788 | 0.166 | 1.000 |

P-values were adjusted for the comparison of 15 genes using Bonferroni procedure. S1-S11 indicates Sample 1-11. *** indicates a $P_{adj}$-value < 0.001.

**Supplementary Table S5. The 12 differentially expressed genes (DEGs) in KO vs. WT samples with $log_2$-fold change (LFC, cutoff = 1) and statistical significance (p-value, corrected for multiple testing) (Fig. 2B-C).**

| | $log_2$ fold change | p-value |
|---|---|---|
| ***Cacna1h*** | -1.348 | $3.9x10^{-05}$ |
| ***Usp2*** | -1.064 | $7.7x10^{-03}$ |
| ***2810039B14Rik*** | -1.058 | $2.4x10^{-04}$ |
| ***Scn8a*** | -1.032 | 0.026 |
| ***Cdv3*** | 1.035 | 0.042 |
| ***2010003K11Rik*** | 1.044 | $9.4x10^{-04}$ |
| ***Ptp4a1*** | 1.055 | $5.4x10^{-04}$ |
| ***Mpp7*** | 1.061 | $5.4x10^{-04}$ |
| ***Creb5*** | 1.068 | $8.4x10^{-03}$ |
| ***Tmprss11a*** | 1.455 | 0.022 |
| ***Gadd45g*** | 1.493 | $4.3x10^{-08}$ |
| ***Lars2*** | 1.82 | 0.036 |

**Supplementary Table S6. Genes of the aldosterone synthesis and secretion pathway (KEGG mmu04925) with log$_2$-fold change (LFC) in gene expression KO vs. WT samples and the statistical significance (p-value, corrected for multiple testing).**

| Symbol | LFC | p-value |
|---|---:|---:|
| Cacna1h | -1,348 | 3,9E-05 |
| Star | 0,347 | 0,007 |
| Creb5 | 1,068 | 0,008 |
| Gna11 | -0,198 | 0,023 |
| Agtr1a | 0,431 | 0,048 |
| Creb1 | 0,193 | 0,067 |
| Scarb1 | 0,309 | 0,118 |
| Dagla | -0,356 | 0,157 |
| Orai1 | -0,170 | 0,237 |
| Kcnj5 | -0,054 | 0,252 |
| Plcb3 | -0,143 | 0,259 |
| Camk1 | -0,111 | 0,268 |
| Nr4a2 | -0,012 | 0,297 |
| Camk2d | 0,080 | 0,403 |
| Prkce | 0,079 | 0,419 |
| Atf6b | -0,077 | 0,479 |
| Hsd3b6 | -0,016 | 0,501 |
| Kcnk3 | 0,068 | 0,509 |
| Camk4 | 0,016 | 0,527 |
| Prkaca | -0,060 | 0,565 |
| Atp1a1 | 0,066 | 0,590 |
| Atp2b1 | 0,043 | 0,613 |
| Lipe | 0,022 | 0,682 |
| Pde2a | -0,019 | 0,695 |
| Plcb2 | -0,019 | 0,703 |
| Daglb | -0,046 | 0,718 |
| Adcy8 | 0,010 | 0,720 |
| Prkd1 | -0,042 | 0,731 |
| Adcy9 | -0,042 | 0,737 |
| Adcy5 | 0,028 | 0,769 |
| Cacna1c | 0,037 | 0,771 |
| Prkcb | -0,023 | 0,796 |
| Calm1 | 0,035 | 0,799 |
| Atp2b3 | -0,026 | 0,806 |

| Symbol | LFC | p-value |
|---|---:|---:|
| Atp1b1 | 0,023 | 0,812 |
| Atp1b2 | -0,013 | 0,816 |
| Prkacb | 0,033 | 0,820 |
| Itpr2 | 0,027 | 0,828 |
| Calml4 | -0,016 | 0,835 |
| Itpr3 | -0,016 | 0,843 |
| Adcy6 | 0,019 | 0,847 |
| Atf4 | 0,020 | 0,855 |
| Calm3 | -0,026 | 0,857 |
| Itpr1 | 0,024 | 0,860 |
| Adcy3 | -0,016 | 0,864 |
| Agtr1b | -0,021 | 0,866 |
| Kcnk9 | -0,007 | 0,870 |
| Gnaq | 0,025 | 0,870 |
| Gnas | 0,024 | 0,885 |
| Mc2r | -0,010 | 0,887 |
| Cacna1g | 0,006 | 0,887 |
| Nr4a1 | -0,003 | 0,888 |
| Atf2 | -0,020 | 0,902 |
| Hsd3b1 | 0,011 | 0,912 |
| Plcb4 | 0,013 | 0,916 |
| Cacna1i | 0,005 | 0,920 |
| Ldlr | 0,012 | 0,923 |
| Plcb1 | 0,017 | 0,923 |
| Adcy2 | 0,008 | 0,926 |
| Prkca | -0,009 | 0,926 |
| Atp1b3 | 0,016 | 0,932 |
| Creb3l1 | 0,006 | 0,934 |
| Npr1 | -0,013 | 0,938 |
| Adcy1 | 0,006 | 0,940 |
| Creb3l2 | 0,014 | 0,941 |
| Camk2b | -0,004 | 0,955 |
| Creb3 | 0,010 | 0,962 |
| Camk2g | -0,006 | 0,969 |

| Symbol | LFC | p-value |
|---|---:|---:|
| Cyp21a1 | -0,004 | 0,971 |
| Atp2b4 | 0,005 | 0,973 |
| Cyp11a1 | -0,008 | 0,974 |
| Creb3l3 | 0,001 | 0,975 |
| Cacna1d | 0,003 | 0,976 |
| Calm2 | -0,006 | 0,981 |
| Prkd3 | -0,008 | 0,982 |
| Atp2b2 | 0,002 | 0,983 |
| Atp1a2 | -0,004 | 0,984 |
| Atp1a3 | 0,002 | 0,985 |
| Adcy4 | -0,003 | 0,985 |
| Atp1a4 | -0,001 | 0,988 |
| Atf1 | -0,003 | 0,989 |
| Prkd2 | -0,002 | 0,989 |
| Adcy7 | 0,002 | 0,989 |
| Camk2a | -0,001 | 0,992 |
| Agt | 0,001 | 0,993 |
| Hsd3b2 | 0,001 | 0,993 |
| Creb3l4 | -0,012 | # |
| Prkcg | -0,009 | # |
| Cacna1s | -0,007 | # |
| Pomc | -0,006 | # |
| Cacna1f | -0,001 | # |
| Atp1b4 | -0,001 | # |
| Camk1g | 0,000 | # |
| Calml3 | 0,000 | # |
| Hsd3b3 | 0,001 | # |
| Hsd3b8 | ‡ | |
| Hsd3b9 | ‡ | |
| Hsd3b4 | ‡ | |
| Hsd3b5 | ‡ | |
| Nppa | ‡ | |
| Calm5 | ‡ | |
| Calm4 | ‡ | |

### Low mean normalized count

‡ No read counts detected across all samples

*Supplementary Table S7. Patch Clamp recordings (Fig. 2D-E).*

| | WT | | | | | | KO | | | | | | | |
|---|---|---|---|---|---|---|---|---|---|---|---|---|---|---|
| | T-type | | | Non-T-type | | | T-type | | | Non-T-type | | | | |
| **U (mV)** | I (pA) | 95% CI (lower limit) | 95% CI (upper limit) | I (pA) | 95% CI (lower limit) | 95% CI (upper limit) | I (pA) | 95% CI (lower limit) | 95% CI (upper limit) | I (pA) | 95% CI (lower limit) | 95% CI (upper limit) | p-value (T-type) | p-value (Non-T-type) |
| **-95** | ‡ | | | ‡ | | | ‡ | | | ‡ | | | | |
| **-85** | ‡ | | | ‡ | | | ‡ | | | ‡ | | | | |
| **-75** | ‡ | | | ‡ | | | ‡ | | | ‡ | | | | |
| **-65** | ‡ | | | ‡ | | | ‡ | | | ‡ | | | | |
| **-55** | -16.666 | -21.208 | -12.938 | -0.167 | -4.473 | 4.009 | -7.379 | -17.447 | 1.281 | -2.133 | -8.399 | 4.204 | # | # |
| **-45** | -34.305 | -53.887 | -15.06 | -0.68 | -10.106 | 8.533 | -13.143 | -22.237 | -6.423 | -4.302 | -6.249 | -2.275 | # | # |
| **-35** | -49.403 | -70.534 | -29.438 | -8.657 | -13.413 | -3.429 | -10.398 | -19.51 | -2.136 | -5.222 | -10.641 | 0.436 | p= 0.003, ** | # |
| **-25** | -48.19 | -71.216 | -25.164 | -18.712 | -26.757 | -11.479 | -7.04 | -15.232 | 1.471 | -14.054 | -21.388 | -8.532 | # | # |
| **-15** | -45.912 | -65.153 | -25.571 | -28.287 | -44.046 | -14.777 | -10.625 | -26.618 | 3.761 | -18.532 | -23.827 | -13.969 | # | p=0.676, ns |
| **-5** | -42.355 | -66.828 | -23.587 | -31.179 | -45.936 | -17.708 | -5.797 | -19.106 | 5.655 | -24.256 | -31.091 | -17.892 | # | # |
| **5** | -31.97 | -56.364 | -11.757 | -38.536 | -54.818 | -23.023 | -8.314 | -24.82 | 8.286 | -32.279 | -43.212 | -22.469 | # | # |
| **15** | -22.767 | -44.874 | -6.406 | -35.955 | -47.483 | -25.199 | -4.256 | -16.812 | 12.339 | -32.894 | -44.991 | -21.546 | # | # |
| **25** | -6.787 | -22.684 | 4.987 | -41.008 | -52.033 | -29.202 | -2.355 | -17.474 | 18.09 | -31.502 | -46.041 | -18.553 | # | # |
| **35** | -12.857 | -26.476 | 0.463 | -30.859 | -41.409 | -22.117 | 5.098 | -12.989 | 25.352 | -36.032 | -45.875 | -27.46 | # | # |
| **45** | 0.47 | -14.112 | 15.473 | -38.188 | -49.185 | -26.974 | -3.948 | -10.338 | 1.431 | -25.785 | -34.461 | -19.547 | # | # |
| **55** | -2.768 | -13.404 | 5.003 | -28.53 | -39.947 | -19.614 | 0.256 | -13.134 | 14.761 | -16.499 | -25.486 | -8.458 | # | # |
| **65** | 6.34 | -13.252 | 25.932 | -28.676 | -42.26 | -15.092 | 1.585 | -5.67 | 8.69 | -19.975 | -27.238 | -12.051 | # | # |
| **75** | 20.756 | -2.993 | 46.881 | -23.328 | -49.651 | -3.028 | 2.52 | -3.674 | 9.088 | -19.361 | -28.721 | -8.995 | # | # |
| n(animals) | 3 | | | | | | 4 | | | | | | | |
| n(cells) | 7 | | | | | | 9 | | | | | | | |

### not tested

‡ No peak current detected

**Supplementary Table S8. Calcium spike activity during perfusion with 15 µmol/l TTA-P2 in 1/s (Fig. 3E)**

| | WT | | | WT control | | | KO | | | KO control | | |
|---|---|---|---|---|---|---|---|---|---|---|---|---|
| bin (min) | Activity | 95% CI (lower limit) | 95% CI (upper limit) | Activity | 95% CI (lower limit) | 95% CI (upper limit) | Activity | 95% CI (lower limit) | 95% CI (upper limit) | Activity | 95% CI (lower limit) | 95% CI (upper limit) |
| -3 | 0.599 | 0.548 | 0.652 | 0.643 | 0.588 | 0.7 | 1.099 | 1 | 1.197 | 1.173 | 1.025 | 1.313 |
| -2.5 | 0.62 | 0.566 | 0.673 | 0.643 | 0.59 | 0.699 | 1.118 | 1.024 | 1.213 | 1.175 | 1.027 | 1.32 |
| -2 | 0.639 | 0.59 | 0.689 | 0.632 | 0.583 | 0.681 | 1.091 | 0.997 | 1.188 | 1.25 | 1.121 | 1.378 |
| -1.5 | 0.672 | 0.626 | 0.719 | 0.655 | 0.608 | 0.704 | 1.141 | 1.056 | 1.226 | 1.231 | 1.096 | 1.366 |
| -1 | 0.657 | 0.612 | 0.704 | 0.638 | 0.588 | 0.686 | 1.106 | 1.02 | 1.193 | 1.267 | 1.142 | 1.393 |
| -0.5 | 0.623 | 0.575 | 0.671 | 0.612 | 0.566 | 0.658 | 1.12 | 1.039 | 1.203 | 1.218 | 1.096 | 1.339 |
| 0 | 0.56 | 0.51 | 0.611 | 0.592 | 0.547 | 0.636 | 1.068 | 0.983 | 1.155 | 1.194 | 1.081 | 1.304 |
| 0.5 | 0.507 | 0.457 | 0.557 | 0.518 | 0.468 | 0.568 | 1.021 | 0.929 | 1.113 | 1.088 | 0.956 | 1.218 |
| 1 | 0.329 | 0.286 | 0.375 | 0.473 | 0.423 | 0.524 | 0.976 | 0.88 | 1.074 | 1.108 | 0.96 | 1.255 |
| 1.5 | 0.22 | 0.18 | 0.262 | 0.452 | 0.404 | 0.504 | 0.981 | 0.887 | 1.077 | 1.094 | 0.943 | 1.245 |
| 2 | 0.215 | 0.174 | 0.258 | 0.45 | 0.4 | 0.501 | 0.931 | 0.836 | 1.028 | 1.112 | 0.965 | 1.259 |
| 2.5 | 0.241 | 0.196 | 0.286 | 0.441 | 0.396 | 0.489 | 0.954 | 0.857 | 1.051 | 1.123 | 0.986 | 1.256 |
| 3 | 0.261 | 0.211 | 0.311 | 0.443 | 0.396 | 0.492 | 0.924 | 0.833 | 1.015 | 1.058 | 0.923 | 1.191 |
| 3.5 | 0.26 | 0.212 | 0.311 | 0.439 | 0.393 | 0.487 | 0.872 | 0.781 | 0.965 | 1.073 | 0.943 | 1.198 |
| 4 | 0.243 | 0.197 | 0.292 | 0.417 | 0.372 | 0.464 | 0.897 | 0.803 | 0.993 | 0.984 | 0.855 | 1.107 |
| 4.5 | 0.225 | 0.18 | 0.271 | 0.444 | 0.397 | 0.491 | 0.829 | 0.737 | 0.922 | 0.945 | 0.798 | 1.09 |
| 5 | 0.231 | 0.186 | 0.28 | 0.446 | 0.402 | 0.491 | 0.885 | 0.797 | 0.973 | 0.963 | 0.831 | 1.098 |
| 5.5 | 0.221 | 0.176 | 0.267 | 0.427 | 0.382 | 0.473 | 0.859 | 0.768 | 0.949 | 0.969 | 0.843 | 1.098 |
| 6 | 0.221 | 0.178 | 0.268 | 0.426 | 0.38 | 0.473 | 0.848 | 0.755 | 0.942 | 0.894 | 0.753 | 1.036 |
| 6.5 | 0.24 | 0.192 | 0.29 | 0.434 | 0.388 | 0.481 | 0.858 | 0.769 | 0.95 | 0.948 | 0.809 | 1.09 |
| 7 | 0.216 | 0.173 | 0.262 | 0.428 | 0.381 | 0.476 | 0.851 | 0.759 | 0.946 | 0.898 | 0.756 | 1.042 |
| 7.5 | 0.201 | 0.158 | 0.245 | 0.361 | 0.316 | 0.408 | 0.852 | 0.76 | 0.944 | 0.855 | 0.725 | 0.992 |

**Supplementary Table S9. Statistical information of calcium bursting parameters relative to control at 15 µmol/l TTA-P2. (Fig. 3F and G)**

| | WT | | | KO | | |
|---|---|---|---|---|---|---|
| | Mean | 95% CI (lower limit) | 95% CI (upper limit) | Mean | 95% CI (lower limit) | 95% CI (upper limit) |
| Activity | 0.569 | 0.341 | 0.823 | 0.995 | 0.851 | 1.164 |
| Intra-burst frequency | 1.182 | 1.101 | 1.265 | 0.913 | 0.837 | 0.995 |

**Supplementary Table S10. Statistical information of absolute calcium bursting parameters at 15 µmol/l TTA-P2 in WT slices (Fig. 3D and S3A-B) and number of recorded animals, slices and cells.**

| | WT | | | WT control | | | |
|---|---|---|---|---|---|---|---|
| | Mean | CI (lower limit) | CI (upper limit) | Mean | CI (lower limit) | CI (upper limit) | p-value |
| Activity (1/s) | 0.256 | 0.153 | 0.370 | 0.449 | 0.348 | 0.597 | 0.003, ** |
| Intra-burst frequency (1/s) | 0.972 | 0.906 | 1.041 | 0.851 | 0.747 | 1.003 | 0.047, * |
| Active cells per slice (%) | 34.371 | 22.233 | 47.316 | 75.494 | 68.198 | 82.513 | 2.513x10-6, ** |
| n(animals) | 10 | | | 15 | | | |
| n(slices) | 16 | | | 17 | | | |
| n(cells) | 291 | | | 288 | | | |

**Supplementary Table S11. Statistical information of absolute calcium bursting parameters at 15 µmol/l TTA-P2 in KO slices (Fig. 3D and S3A, C) and number of recorded animals, slices and cells.**

| | KO | | | KO control | | | |
|---|---|---|---|---|---|---|---|
| | Mean | CI (lower limit) | CI (upper limit) | Mean | CI (lower limit) | CI (upper limit) | p-value |
| Activity (1/s) | 0.871 | 0.745 | 1.018 | 0.875 | 0.662 | 1.096 | 0.958, ns |
| Intra-burst frequency (1/s) | 1.239 | 1.136 | 1.351 | 1.307 | 1.162 | 1.459 | 0.446, ns |
| Active cells per slice (%) | 90.288 | 84.286 | 95.110 | 83.588 | 73.129 | 93.793 | 0.230, ns |
| n(animals) | 7 | | | 7 | | | |
| n(slices) | 13 | | | 7 | | | |
| n(cells) | 162 | | | 83 | | | |

**Supplementary Table S12. Statistical information of absolute intra-burst frequency (1/s) at 0 µmol/l TTA-P2 in remaining active WT, KO and corresponding control slices (Fig. S3B-C).**

| | WT | | | WT control | | | |
|---|---|---|---|---|---|---|---|
| | Mean | CI (lower limit) | CI (upper limit) | Mean | CI (lower limit) | CI (upper limit) | p-value |
| Intra-burst frequency (1/s) | 0.929 | 0.840 | 1.029 | 0.960 | 0.841 | 1.114 | 0.9769, ns |
| | KO | | | KO control | | | |
| Intra-burst frequency (1/s) | 1.307 | 1.196 | 1.444 | 1.410 | 1.291 | 1.533 | 0.2702, ns |

**Supplementary Table S13. Calcium spike activity during perfusion with 0.05 µmol/l isradipine in 1/s (Fig. 4C)**

| | WT | | | WT control | | | KO | | | KO control | | |
|---|---|---|---|---|---|---|---|---|---|---|---|---|
| bin (min) | Activity | 95% CI (lower limit) | 95% CI (upper limit) | Activity | 95% CI (lower limit) | 95% CI (upper limit) | Activity | 95% CI (lower limit) | 95% CI (upper limit) | Activity | 95% CI (lower limit) | 95% CI (upper limit) |
| -3 | 0.544 | 0.474 | 0.617 | 0.643 | 0.588 | 0.7 | 1.223 | 1.105 | 1.346 | 1.173 | 1.025 | 1.313 |
| -2.5 | 0.559 | 0.487 | 0.632 | 0.643 | 0.59 | 0.699 | 1.304 | 1.195 | 1.418 | 1.175 | 1.027 | 1.32 |
| -2 | 0.597 | 0.527 | 0.669 | 0.632 | 0.583 | 0.681 | 1.314 | 1.197 | 1.431 | 1.25 | 1.121 | 1.378 |
| -1.5 | 0.649 | 0.582 | 0.719 | 0.655 | 0.608 | 0.704 | 1.278 | 1.157 | 1.4 | 1.231 | 1.096 | 1.366 |
| -1 | 0.63 | 0.564 | 0.698 | 0.638 | 0.588 | 0.686 | 1.253 | 1.137 | 1.371 | 1.267 | 1.142 | 1.393 |
| -0.5 | 0.562 | 0.496 | 0.631 | 0.612 | 0.566 | 0.658 | 1.213 | 1.087 | 1.34 | 1.218 | 1.096 | 1.339 |
| 0 | 0.563 | 0.503 | 0.625 | 0.592 | 0.547 | 0.636 | 1.194 | 1.067 | 1.321 | 1.194 | 1.081 | 1.304 |
| 0.5 | 0.53 | 0.467 | 0.596 | 0.518 | 0.468 | 0.568 | 1.207 | 1.085 | 1.33 | 1.088 | 0.956 | 1.218 |
| 1 | 0.492 | 0.424 | 0.563 | 0.473 | 0.423 | 0.524 | 1.086 | 0.962 | 1.209 | 1.108 | 0.96 | 1.255 |
| 1.5 | 0.493 | 0.433 | 0.556 | 0.452 | 0.404 | 0.504 | 0.887 | 0.767 | 1.009 | 1.094 | 0.943 | 1.245 |
| 2 | 0.468 | 0.405 | 0.536 | 0.45 | 0.4 | 0.501 | 0.872 | 0.749 | 0.999 | 1.112 | 0.965 | 1.259 |
| 2.5 | 0.375 | 0.315 | 0.437 | 0.441 | 0.396 | 0.489 | 0.884 | 0.766 | 1.005 | 1.123 | 0.986 | 1.256 |
| 3 | 0.342 | 0.284 | 0.402 | 0.443 | 0.396 | 0.492 | 0.799 | 0.678 | 0.925 | 1.058 | 0.923 | 1.191 |
| 3.5 | 0.368 | 0.311 | 0.428 | 0.439 | 0.393 | 0.487 | 0.614 | 0.509 | 0.72 | 1.073 | 0.943 | 1.198 |
| 4 | 0.382 | 0.323 | 0.443 | 0.417 | 0.372 | 0.464 | 0.505 | 0.402 | 0.613 | 0.984 | 0.855 | 1.107 |
| 4.5 | 0.388 | 0.327 | 0.451 | 0.444 | 0.397 | 0.491 | 0.457 | 0.362 | 0.559 | 0.945 | 0.798 | 1.09 |
| 5 | 0.371 | 0.31 | 0.433 | 0.446 | 0.402 | 0.491 | 0.316 | 0.231 | 0.408 | 0.963 | 0.831 | 1.098 |
| 5.5 | 0.361 | 0.298 | 0.424 | 0.427 | 0.382 | 0.473 | 0.271 | 0.191 | 0.356 | 0.969 | 0.843 | 1.098 |
| 6 | 0.343 | 0.285 | 0.404 | 0.426 | 0.38 | 0.473 | 0.198 | 0.131 | 0.27 | 0.894 | 0.753 | 1.036 |
| 6.5 | 0.361 | 0.308 | 0.415 | 0.434 | 0.388 | 0.481 | 0.218 | 0.147 | 0.292 | 0.948 | 0.809 | 1.09 |
| 7 | 0.373 | 0.314 | 0.435 | 0.428 | 0.381 | 0.476 | 0.141 | 0.087 | 0.203 | 0.898 | 0.756 | 1.042 |
| 7.5 | 0.387 | 0.324 | 0.452 | 0.361 | 0.316 | 0.408 | 0.115 | 0.064 | 0.175 | 0.855 | 0.725 | 0.992 |

**Supplementary Table S14. Statistical information of absolute calcium bursting parameters at 0.05 µmol/l isradipine in WT slices (Fig. S4A-B)**

| | WT | | | WT control | | | |
|---|---|---|---|---|---|---|---|
| | Mean | 95% CI (lower limit) | 95% CI (upper limit) | Mean | 95% CI (lower limit) | 95% CI (upper limit) | p-value |
| Activity (1/s) | 0.350 | 0.264 | 0.444 | 0.449 | 0.348 | 0.597 | 0.255, ns |
| Intra-burst frequency (1/s) | 0.838 | 0.656 | 1.023 | 0.851 | 0.747 | 1.003 | 0.749, ns |
| n(animals) | 9 | | | 15 | | | |
| n(slices) | 10 | | | 17 | | | |
| n(cells) | 178 | | | 288 | | | |

**Supplementary Table S15. Statistical information of absolute calcium bursting parameters at 0.05 µmol/l isradipine in KO slices (Fig. S4A, S4C)**

| | KO | | | KO control | | | |
|---|---|---|---|---|---|---|---|
| | Mean | 95% CI (lower limit) | 95% CI (upper limit) | Mean | 95% CI (lower limit) | 95% CI (upper limit) | p-value |
| Activity (1/s) | 0.144 | 0.088 | 0.217 | 0.875 | 0.662 | 1.096 | $3.107 \times 10^{-6}$, *** |
| Intra-burst frequency (1/s) | 1.312 | 1.241 | 1.382 | 1.307 | 1.162 | 1.459 | 0.879, ns |
| n(animals) | 6 | | | 7 | | | |
| n(slices) | 7 | | | 7 | | | |
| n(cells) | 107 | | | 83 | | | |

**Supplementary Table S16. Statistical information of absolute intra-burst frequency (1/s) at 0 µmol/l isradipine in remaining active WT, KO and corresponding control slices (Fig. S4B-C)**

| | WT | | | WT control | | | |
|---|---|---|---|---|---|---|---|
| | Mean | CI (lower limit) | CI (upper limit) | Mean | CI (lower limit) | CI (upper limit) | p-value |
| Intra-burst frequency (1/s) | 0.868 | 0.670 | 1.078 | 0.960 | 0.841 | 1.114 | 0.2939, ns |
| | KO | | | KO control | | | |
| Intra-burst frequency (1/s) | 1.363 | 1.218 | 1.507 | 1.410 | 1.291 | 1.533 | 0.4994, ns |

**Supplementary Table S17. Calcium spike activity during perfusion with 5 µmol/l TTA-P2 and 300 nmol/l isradipine in 1/s (Fig. 5B)**

| | WT | | | WT control | | |
|---|---|---|---|---|---|---|
| Bins (min) | Activity | 95 % CI (lower limit) | 95 % CI (upper limit) | Activity | 95 % CI (lower limit) | 95 % CI (upper limit) |
| -4 | 0.619 | 0.578 | 0.661 | 0.697 | 0.645 | 0.747 |
| -3 | 0.571 | 0.536 | 0.607 | 0.674 | 0.62 | 0.729 |
| -2 | 0.522 | 0.486 | 0.561 | 0.642 | 0.591 | 0.695 |
| -1 | 0.545 | 0.511 | 0.579 | 0.624 | 0.577 | 0.672 |
| 0 | 0.518 | 0.485 | 0.553 | 0.58 | 0.539 | 0.62 |
| 1 | 0.439 | 0.399 | 0.481 | 0.472 | 0.425 | 0.521 |
| 2 | 0.282 | 0.241 | 0.324 | 0.443 | 0.398 | 0.491 |
| 3 | 0.226 | 0.189 | 0.264 | 0.44 | 0.397 | 0.483 |
| 4 | 0.199 | 0.159 | 0.242 | 0.4 | 0.358 | 0.443 |
| 5 | 0.17 | 0.131 | 0.211 | 0.414 | 0.371 | 0.459 |
| 6 | 0.155 | 0.119 | 0.192 | 0.39 | 0.347 | 0.434 |
| 7 | 0.135 | 0.101 | 0.169 | 0.405 | 0.36 | 0.453 |
| 8 | 0.146 | 0.108 | 0.185 | 0.321 | 0.277 | 0.366 |
| 9 | 0.097 | 0.067 | 0.129 | 0.325 | 0.283 | 0.368 |
| 10 | 0.104 | 0.075 | 0.136 | 0.314 | 0.274 | 0.354 |
| 11 | 0.097 | 0.067 | 0.131 | 0.315 | 0.274 | 0.355 |
| 12 | 0.063 | 0.04 | 0.089 | 0.254 | 0.216 | 0.294 |
| 13 | 0.039 | 0.022 | 0.059 | 0.301 | 0.258 | 0.344 |
| 14 | 0.013 | 0.004 | 0.023 | 0.241 | 0.203 | 0.279 |
| 15 | 0.003 | 0 | 0.006 | 0.256 | 0.221 | 0.293 |
| 16 | 0 | 0 | 0 | 0.221 | 0.186 | 0.257 |
| 17 | 0 | 0 | 0.001 | 0.225 | 0.191 | 0.261 |
| 18 | 0 | 0 | 0 | 0.21 | 0.175 | 0.246 |
| 19 | 0 | 0 | 0 | 0.218 | 0.181 | 0.256 |
| 20 | 0 | 0 | 0 | 0.201 | 0.165 | 0.237 |

**Supplementary Table S18. Statistical information of calcium bursting parameters relative to control in WT slices (Fig. 5C and S5C)**

| | 5 µmol/l TTA-P2 | | | 5 µmol/l TTA-P2 and 300 nmol/l isradipine | | |
|---|---|---|---|---|---|---|
| | Mean | 95% CI (lower limit) | 95% CI (upper limit) | Mean | 95% CI (lower limit) | 95% CI (upper limit) |
| Activity | 0.250 | 0.12 | 0.402 | 0.001 | 0 | 0.002 |
| Intra-burst frequency | 1.341 | 1.137 | 1.536 | --- | --- | --- |

**Supplementary Table S19. Statistical information of absolute calcium bursting parameters at 5 µmol/l TTA-P2 in WT slices (Fig. S5A, B, D)**

| | WT | | | WT control | | | |
|---|---|---|---|---|---|---|---|
| | Mean | 95% CI (lower limit) | 95% CI (upper limit) | Mean | 95% CI (lower limit) | 95% CI (upper limit) | p-value |
| Activity (1/s) | 0.078 | 0.037 | 0.125 | 0.310 | 0.257 | 0.362 | $3.248 \times 10^{-05}$, *** |
| Intra-burst frequency (1/s) | 1.022 | 0.867 | 1.171 | 0.762 | 0.685 | 0.848 | 0.006, ** |
| Active cells per slice (%) | 16.611 | 8.989 | 25.094 | 63.003 | 52.350 | 73.262 | $2.371 \times 10^{-07}$, *** |
| n(animals) | 6 | | | 9 | | | |
| n(slices) | 10 | | | 10 | | | |
| n(cells) | 246 | | | 213 | | | |

**Supplementary Table S20. Statistical information of absolute calcium bursting parameters at 5 µmol/l TTA-P2 and 300 nmol/l isradipine in WT slices (Fig. S5A, B)**

| | WT | | | WT control | | | |
|---|---|---|---|---|---|---|---|
| | Mean | 95% CI (lower limit) | 95% CI (upper limit) | Mean | 95% CI (lower limit) | 95% CI (upper limit) | p-value |
| Activity (1/s) | 0 | 0 | 0 | 0.208 | 0.153 | 0.262 | $2.839 \times 10^{-13}$, *** |
| Active cells per slice (%) | 0 | 0 | 0 | 48.541 | 35.409 | 62.104 | $5.572 \times 10^{-09}$, *** |

**Supplementary Table S21. Statistical information of absolute intra-burst frequency of remaining active cells at 0 µmol/l TTA-P2 and isradipine in WT and WT control slices (Fig. S5D)**

| | WT | | | WT control | | | |
|---|---|---|---|---|---|---|---|
| | Mean | 95% CI (lower limit) | 95% CI (upper limit) | Mean | 95% CI (lower limit) | 95% CI (upper limit) | p-value |
| Intra-burst frequency (1/s) | 0.651 | 0.572 | 0.755 | 0.896 | 0.782 | 1.034 | 0.018, * |

**Supplementary Table S22. Mean $[Ca^{2+}]_i$ during perfusion with 5 µmol/l TTA-P2 and 300 nmol/l isradipine at various Ang II concentrations in nmol/l in WT cells and WT control (no inhibitors) (Fig. 5D).**

| | WT | | | WT control | | |
|---|---|---|---|---|---|---|
| Bins (min) | $[Ca^{2+}]_i$ | 95 % CI (lower limit) | 95 % CI (upper limit) | $[Ca^{2+}]_i$ | 95 % CI (lower limit) | 95 % CI (upper limit) |
| -4 | 864.549 | 836.929 | 892.926 | 888.099 | 854.841 | 922.111 |
| -3 | 865.652 | 838.906 | 893.437 | 891.308 | 858.863 | 924.363 |
| -2 | 869.298 | 841.805 | 897.773 | 885.136 | 853.188 | 916.859 |
| -1 | 868.608 | 840.997 | 896.654 | 898.190 | 865.840 | 930.566 |
| 0 | 867.532 | 840.128 | 895.610 | 898.049 | 866.559 | 929.423 |
| 1 | 870.386 | 842.817 | 898.588 | 930.176 | 894.131 | 966.837 |
| 2 | 902.025 | 871.246 | 933.615 | 983.125 | 939.729 | 1027.534 |
| 3 | 954.147 | 917.252 | 991.973 | 1000.043 | 960.741 | 1040.524 |
| 4 | 977.727 | 940.114 | 1016.438 | 1032.237 | 988.984 | 1076.662 |
| 5 | 983.641 | 946.009 | 1022.706 | 1036.660 | 991.294 | 1082.005 |
| 6 | 973.729 | 939.878 | 1008.134 | 1108.969 | 1058.135 | 1159.981 |
| 7 | 971.776 | 939.103 | 1005.421 | 1058.114 | 1011.845 | 1107.226 |
| 8 | 998.934 | 960.843 | 1037.959 | 1036.406 | 993.498 | 1080.996 |
| 9 | 983.862 | 943.155 | 1026.580 | 1043.866 | 1004.186 | 1085.725 |
| 10 | 1006.429 | 966.100 | 1049.140 | 1023.154 | 978.285 | 1070.870 |
| 11 | 973.993 | 938.483 | 1010.298 | 1027.249 | 981.977 | 1075.130 |
| 12 | 909.260 | 881.889 | 936.394 | 1034.765 | 989.021 | 1083.589 |
| 13 | 871.519 | 845.934 | 897.578 | 980.283 | 943.726 | 1017.967 |
| 14 | 862.802 | 836.559 | 888.990 | 998.539 | 963.245 | 1035.649 |
| 15 | 850.449 | 824.585 | 876.181 | 1000.975 | 964.852 | 1038.451 |
| 16 | 846.927 | 821.062 | 872.230 | 975.663 | 935.696 | 1017.947 |
| 17 | 841.812 | 817.417 | 865.882 | 996.340 | 954.660 | 1038.854 |
| 18 | 843.909 | 819.944 | 867.792 | 1000.799 | 963.758 | 1040.706 |
| 19 | 842.062 | 818.451 | 865.552 | 953.245 | 917.400 | 989.815 |
| 20 | 838.415 | 815.881 | 861.054 | 971.624 | 933.894 | 1010.200 |
| 21 | 832.643 | 810.607 | 855.098 | 986.278 | 948.122 | 1024.644 |
| 22 | 847.909 | 824.713 | 871.310 | 1046.175 | 1008.910 | 1083.971 |
| 23 | 854.322 | 831.536 | 877.100 | 1050.743 | 1010.929 | 1090.409 |
| 24 | 852.572 | 830.638 | 874.356 | 1053.052 | 1016.071 | 1090.860 |
| 25 | 842.792 | 821.377 | 863.782 | 1099.930 | 1063.676 | 1136.284 |
| 26 | 834.369 | 813.755 | 855.144 | 1079.310 | 1043.213 | 1115.221 |
| 27 | 823.902 | 804.253 | 843.816 | 1067.017 | 1030.410 | 1103.287 |
| 28 | 817.811 | 798.443 | 837.846 | 1075.528 | 1042.072 | 1109.497 |
| 29 | 816.627 | 798.123 | 835.755 | 1069.909 | 1037.969 | 1102.267 |
| 30 | 793.104 | 765.087 | 824.458 | 1057.602 | 1024.909 | 1090.494 |

**Supplementary Table S23. Mean baseline $[Ca^{2+}]_i$ during perfusion with 5 µmol/l TTA-P2 and 300 nmol/l isradipine at various Ang II concentrations in nmol/l in WT cells and WT control (no inhibitors) (Fig. 5D).**

| | WT | | | WT control | | |
|---|---|---|---|---|---|---|
| Bins (min) | Baseline $[Ca^{2+}]_i$ | 95 % CI (lower limit) | 95 % CI (upper limit) | Baseline $[Ca^{2+}]_i$ | 95 % CI (lower limit) | 95 % CI (upper limit) |
| -4 | 859.876 | 831.604 | 888.505 | 873.027 | 841.843 | 903.763 |
| -3 | 859.027 | 832.326 | 885.803 | 881.753 | 851.764 | 912.312 |
| -2 | 861.425 | 834.133 | 889.481 | 881.066 | 848.904 | 912.596 |
| -1 | 858.330 | 831.529 | 885.791 | 892.199 | 860.664 | 923.614 |
| 0 | 857.752 | 830.754 | 885.190 | 891.762 | 860.330 | 922.837 |
| 1 | 862.609 | 834.929 | 891.073 | 911.551 | 877.542 | 944.952 |
| 2 | 879.763 | 850.252 | 910.233 | 939.589 | 900.227 | 978.250 |
| 3 | 900.661 | 867.702 | 934.623 | 949.212 | 914.233 | 985.023 |
| 4 | 897.787 | 865.578 | 930.691 | 950.418 | 912.135 | 987.185 |
| 5 | 905.287 | 872.870 | 938.693 | 943.250 | 908.057 | 978.630 |
| 6 | 908.940 | 876.583 | 941.743 | 946.982 | 905.857 | 987.722 |
| 7 | 908.073 | 875.425 | 941.506 | 947.290 | 910.585 | 984.288 |
| 8 | 890.809 | 859.109 | 923.233 | 929.640 | 893.946 | 965.165 |
| 9 | 881.171 | 851.287 | 912.028 | 928.225 | 894.580 | 962.347 |
| 10 | 885.196 | 852.928 | 917.900 | 930.274 | 895.200 | 965.148 |
| 11 | 895.400 | 865.839 | 925.796 | 935.826 | 900.910 | 969.836 |
| 12 | 875.110 | 847.987 | 902.721 | 923.621 | 890.759 | 955.517 |
| 13 | 862.243 | 836.134 | 888.658 | 920.755 | 887.167 | 954.410 |
| 14 | 856.032 | 829.443 | 882.551 | 922.507 | 891.196 | 954.264 |
| 15 | 848.399 | 822.093 | 874.735 | 930.808 | 897.592 | 964.449 |
| 16 | 846.078 | 820.242 | 871.507 | 924.076 | 891.323 | 957.127 |
| 17 | 841.812 | 817.417 | 865.882 | 922.972 | 889.397 | 956.285 |
| 18 | 843.909 | 819.944 | 867.792 | 930.758 | 899.708 | 961.392 |
| 19 | 842.062 | 818.451 | 865.552 | 926.340 | 893.172 | 960.548 |
| 20 | 838.415 | 815.881 | 861.054 | 931.187 | 895.798 | 966.774 |

**Supplementary Table S24. Absolute mean $[Ca^{2+}]_i$ during perfusion with 5 µmol/l TTA-P2 and 300 nmol/l isradipine at various Ang II concentrations in WT cells compared to control (no blocker, Fig. S6A).**

| | | WT | | | WT control (no blocker) | | | |
|---|---|---|---|---|---|---|---|---|
| Ang II (pmol/l) | **Blocker** | $[Ca2+]_i$ | 95 % CI (lower limit) | 95 % CI (upper limit) | $[Ca2+]_i$ | 95 % CI (lower limit) | 95 % CI (upper limit) | p-value |
| 0 | **-** | 841.533 | 747.795 | 915.955 | 903.262 | 885.679 | 925.200 | 0.274, ns |
| 100 | **-** | 951.938 | 822.231 | 1077.177 | 1024.507 | 967.314 | 1096.333 | 0.391, ns |
| 100 | **+** | 817.096 | 733.205 | 900.988 | 968.964 | 917.150 | 1000.945 | 0.015, * |
| n(animals) | | 5 | | | 3 | | | |
| n(slices) | | 5 | | | 4 | | | |
| n(cells) | | 117 | | | 69 | | | |

**Supplementary Table S25. Percentage of time spent bursting during perfusion with 5 µmol/l TTA-P2 and 300 nmol/l isradipine at various Ang II concentrations in WT and untreated control cells (no blocker, Fig. S6B).**

| | | WT | | | WT control (no blocker) | | |
|---|---|---|---|---|---|---|---|
| Ang II (pmol/l) | **Blocker** | % | 95 % CI (lower limit) | 95 % CI (upper limit) | % | 95 % CI (lower limit) | 95 % CI (upper limit) |
| 0 | - | 5.953 | 2.642 | 9.800 | 4.101 | 1.382 | 7.683 |
| 100 | - | 44.323 | 40.082 | 48.493 | 39.547374 | 33.858 | 45.371 |
| 100 | + | - | - | - | 27.025 | 20.822 | 33.775 |
